# A Benchmark Dataset for Machine Learning in Ecotoxicology

**DOI:** 10.1101/2023.05.27.542160

**Authors:** Christoph Schür, Lilian Gasser, Fernando Perez-Cruz, Kristin Schirmer, Marco Baity-Jesi

## Abstract

The use of machine learning for predicting ecotoxicological outcomes is promising, but underutilized. The curation of data with informative features requires both expertise in machine learning as well as a strong biological and ecotoxicological background, which we consider a barrier of entry for this kind of research. Additionally, model performances can only be compared across studies when the same dataset, cleaning, and splittings were used. Therefore, we provide *ADORE*, an extensive and well-described dataset on acute aquatic toxicity in three relevant taxonomic groups (fish, crustaceans, and algae). The core dataset describes ecotoxicological experiments and is expanded with phylogenetic and species-specific data on the species as well as chemical properties and molecular representations. Apart from challenging other researchers to try and achieve the best model performances across the whole dataset, we propose specific relevant challenges on subsets of the data and include datasets and splittings corresponding to each of these challenge as well as in-depth characterization and discussion of train-test splitting approaches.

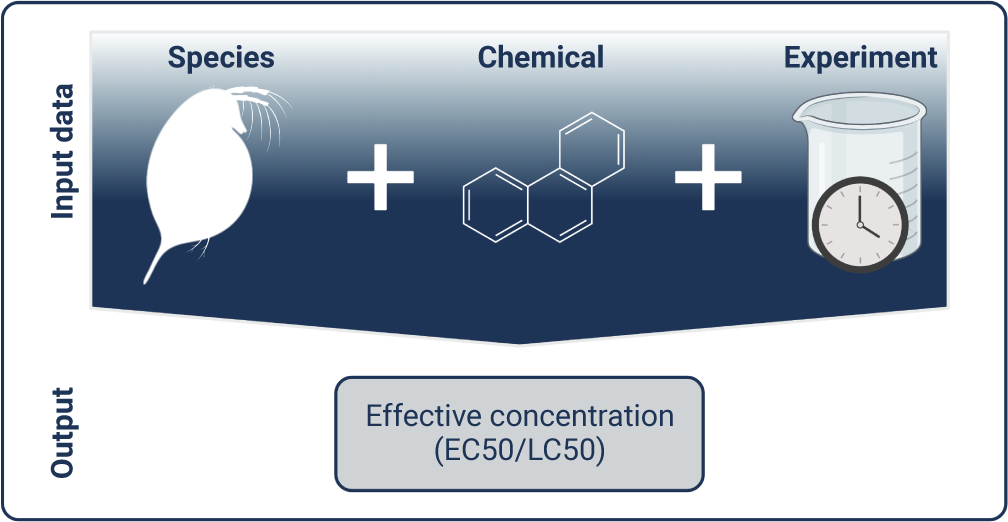

## Background & Summary

Regulation of chemicals aims to protect both human health and the environment, with ecotoxicological research focusing on the latter. Regulatory hazard assessment relies on extensive animal testing, *e.g.,* in the European Union, acute fish toxicity testing for chemicals of a manufacturing or import volume *>* 10 tons p.a. is required by the Registration, Evaluation, Authorisation and Restriction of Chemicals (REACH) legislation^1^. A recent pre-print estimates the global annual usage of fish and birds to range between 440,000 and 2.2 million individuals at a cost upwards of $39 million p.a.^2^. With over 200 million substances in the archive of the Chemical Abstracts Service (CAS, www.cas.org) and more than 350,000 chemicals and mixtures currently registered on the market worldwide^3^, chemical hazard assessment is a major challenge.

These ethical and financial considerations are incentives for alternatives to animal testing, which include computational (*in silico*) methods. A long established *in silico* method is Quantitative structure-activity relationship (QSAR) modeling. It aims to predict biological and chemical properties of compounds based on linear or nonlinear relationships of their molecular structure and experimentally determined activities and properties^4^. The increase of computational capacities and accessibility of educational resources and software led to the adoption of machine learning (ML, computational methods that continually improve their prediction without additional outside intervention) in ecotoxicology over the past decade. Published studies on predicting ecotoxicity are for the most part limited to aquatic systems. They are using animal-derived (*in vivo*) data and encompass supervised learning approaches: classification of chemicals into toxicity brackets, and regression-based prediction of concrete toxicity values^5–7^. The latter more closely resembles the outcome of ecotoxicity experiments, where specimens are exposed to multiple concentrations of the same substance and an effect is recorded at certain time intervals. A non-linear regression model is then fit to the data to derive toxicity values.

Biological and ecotoxicological expertise is required to guide informative feature selection and data cleaning prior to ML modeling, and to help interpret results^8^. In ML-based research, insufficient reporting and data availability hinders reproducibility and comparison across studies and, thus, objective judgment of model performances^9^. Additionally, the way a dataset is split into training and test data immensely affects model performance and can inflate it through data leakage (*i.e.,* the model is trained and tested on similar data)^10^.

Ultimately, realistically assessing and comparing model performance among different works is only feasible when training and testing were performed on the same dataset, using the same data cleaning and splitting strategy. This is fundamental to ML-based research and demonstrated by the plethora of publications associated with long-standing benchmark datasets, such as CIFAR and ImageNet and many others (for example, in the NeurIPS benchmark data track). Compared to NeurIPS, environmental sciences are slowly progressing towards adopting the best practices in the field of ML, including the availability of standard datasets, with a notable pioneer being the CAMELS dataset from hydrology^11,12^ with its recent evolution to CARAVAN^13,14^. Indeed, to serve as benchmark requires a well-characterized, expert-curated and -described, freely available dataset, which we provide with the *ADORE* dataset in this work. The focus of our dataset is mortality and related endpoints for the three taxonomic groups fish, crustaceans, and algae, extracted from the ECOTOX database^15^. The dataset is supplemented by chemical, phylogenetic, and species-specific features that can be used in modeling toxicity outcomes or in addressing other questions.

Here, we describe the data sources and the reasoning behind data selection and processing steps. We provide defined splits of the data, based on chemical occurrence and molecular scaffolds. We propose concrete challenges to the community, including extrapolation across taxonomic groups, to learn more about the potential and limitations of ML in ecotoxicology.

## Methods

The dataset is assembled from different sources. The core contains ecotoxicology data, *e.g.,* toxicity values and experimental conditions. These are expanded with species-related and chemical data (Figure 1). Data sources, filtering and other processing steps are described in the following subsections. The section “Data records” is focused on the different files that contain subsets of the data, whereas data characteristics are shown in the Section “Technical validation”. A glossary of all features is given in Annex Table 6.

**Figure 1.**
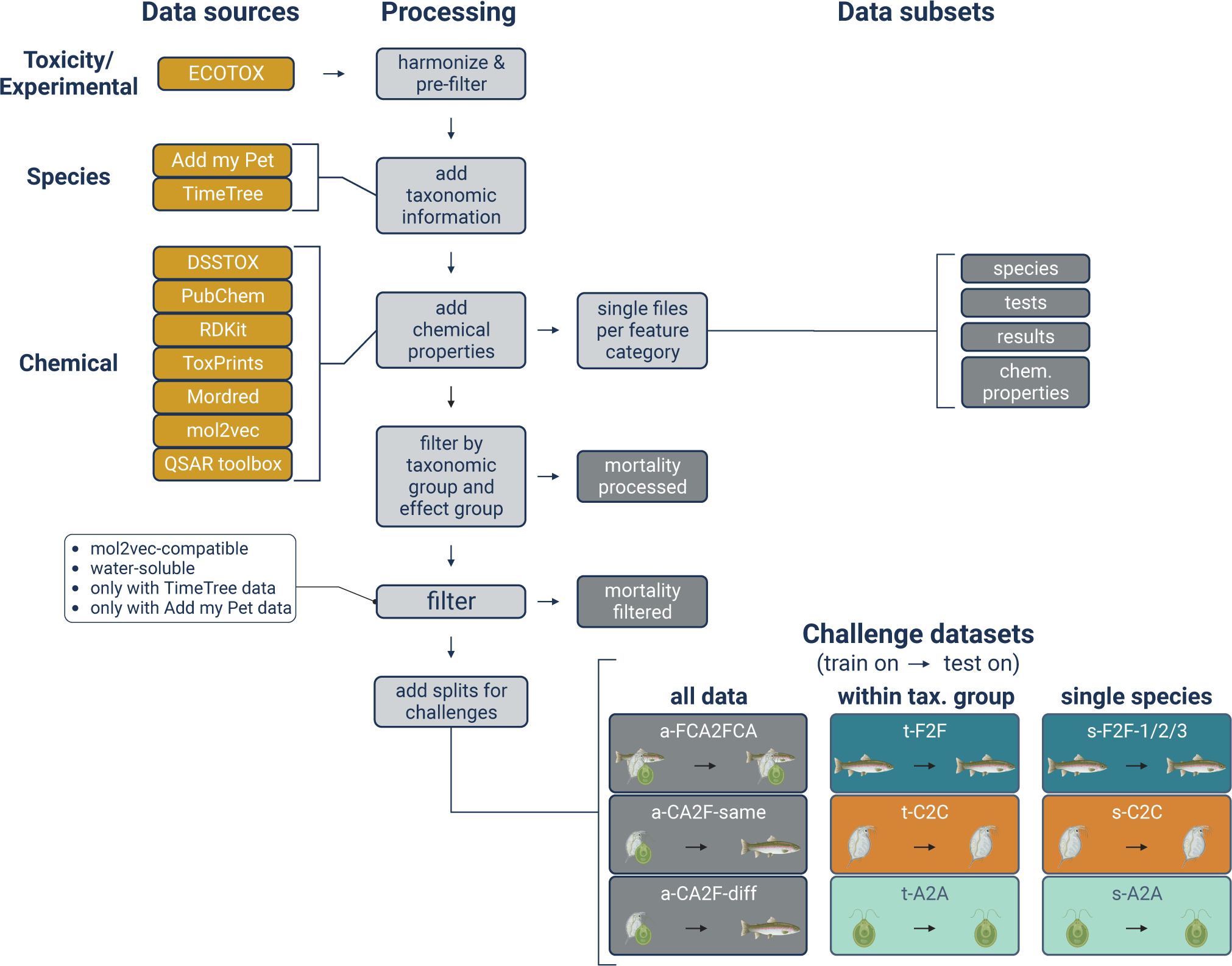
Conceptual overview of the data sources and processing steps.

### Ecotoxicology data

#### Core dataset on acute mortality

The main source of our dataset is the ECOTOX database (https://cfpub.epa.gov/ecotox/) from the United States Environmental Protection Agency’s (US EPA). ECOTOX is updated on a quarterly basis. Our dataset is based on the release from September 2022, which contains over 1.1 million entries of more than 12,000 chemicals and close to 14,000 species^15^. The EnviroTox database^16^ is similar to our dataset, but without the additional chemical- and species-specific features which we curated with ML modeling in mind, and without a standardization that allows straightforward comparability between studies. We had to evaluate a trade-off between a more noisy dataset encompassing more chemical and organismal diversity, and a substantially smaller, but cleaner dataset that is not as chemically and taxonomically diverse.

We include entries for three distinct taxonomic and trophic levels of importance in aquatic ecotoxicology: fish, crustaceans, and algae. These taxonomic groups represent some of the biggest data subsets available in ECOTOX by covering 41% of all the entries.

ECOTOX distinguishes between effects and endpoints. Effects are observed responses resulting from chemical exposure, ranging from mortality (MOR), *i.e.,* death of the specimen, to more subtle effects like behavioral changes, and gene regulation. Endpoints are the targets which can be numerically determined, *e.g.,*, LC50 and EC50. The use of the lethal concentration 50 (LC50), the concentration at which 50% of specimen died after a certain time period, is very common and allows for comparison of toxicity across chemicals and species. In aquatic ecotoxicology, the LC50 refers to the concentration of a substance in the test medium water surrounding a species^17^. For effects that are not lethality-based, toxicity is commonly described as the effective concentration 50 (EC50), that analogously describes the induction of a 50% effect level, often compared to a negative and/or positive control treatment. Both can be expressed in a mass-based (*e.g., mg/L*) and a mol-based (*e.g., mol/L*) manner. The former is easier to correlate to environmental concentrations, while the latter is more biologically informative, assuming that a molecule is the entity that induces the biological response on a molecular level.

Our dataset focuses on short-term lethal (acute) mortality and we included other endpoints that are comparable to mortality (Table 1). Effects manifest differently in different taxa. Mortality was the sole effect chosen in fish, based on standardized test guidelines such as the Organisation for Economic Co-operation and Development (OECD) Test guideline 203^18^. For crustaceans, mortality and immobilization, the latter categorized in ECOTOX under intoxication (ITX), are used interchangeably in the respective test guidelines, *e.g.,* OECD Test guideline 202^19^. For algae, the focus is rarely on the individual, but often related to population health/size as a proxy for toxicity. Population growth is the most commonly used approach to testing effects in algae, *e.g.,* OECD guideline 201^20^), which led to the inclusion of data related to effects encoded as mortality, growth (GRO), population (POP), and physiology (PHY) (Table 1).

**Table 1.** Effect groups for different taxonomic groups. Definitions are given in Annex B.

| Fish | Crustaceans | Algae |
| --- | --- | --- |
| Mortality (MOR) | Mortality (MOR) | Mortality (MOR) |
| - | Intoxication (ITX; immobilization) | Growth (GRO) |
| - | - | Population (POP) |
| - | - | Physiology (PHY) |

The standard observational period for fish acute mortality is 96 hours according to OECD guideline 203^18^, where mortality is recorded every 24 hours. For crustaceans, the typical experimental period is 48 hours, *e.g.,* given in OECD guideline 202^19^, while algae are commonly exposed over 72 hours^20^. Only experimental periods up to 96 hours were included.

Since, ultimately, the goal would be to replace animal testing, we excluded *in vitro* endpoints, *i.e.,* assays based on isolated cells^21^, and assays based on early organismal life stages, such as eggs and embryos^22^. They comprise alternative test methods to animal testing^23^. Removing eggs and embryo tests is done under the limitation that most of the entries have uninformative values for the organism life stage and that life stages for the three taxonomic groups are hardly comparable (Annex Figure 11E).

#### Processing of the ECOTOX database

The core dataset was assembled from the downloadable, pipe-delimited ASCII version of the ECOTOX database. It is made available as a set of.txt files. We used the files for species, tests, results and media that contain information on the taxonomy of species, experimental-design-related factors, experimental outcomes, and media characteristics, respectively. The entries of these files can be matched to each other using different unique keys, *e.g.,* the test species is encoded with the key *species_number* that can be used to match the taxonomic information with the test table. Each data point is uniquely identified by the *result_id*.

Chemicals are identified through different descriptors, most prominently the CAS number, the DSSTox Substance ID (DTXSID), associated with the CompTox Chemicals Dashboard by the US EPA, and the IUPAC International Chemical Identifier (InChI) and the associated InChIkey by the International Union of Pure and Applied Chemistry^24^. These identifiers are retained in the dataset to ensure high compatibility with other data sources and retraceability of all data points. If possible, the InChIkey was used for matching, followed by DTXSID and CAS number. Additionally, the dataset contains Simplified Molecular Input Line Entry System (SMILES) codes that represent the chemical structure in a string format^25^. Like others^16,26,27^, we use the canonical SMILES, from PubChem^28,29^, which do not capture isomerism.

In the first step of the processing pipeline (Figure 1), the ECOTOX files were separately harmonized and pre-filtered.

##### Species file

The species file contains information on the test species. We removed all data points that had missing values for the species group (column `ecotox_group`) or the taxonomic columns class, order, family, species and genus. We then retained only entries for the three taxonomic groups of interest. This was done by filtering the species group entries to contain either “Fish”, “Crusta”, or “Algae”.

##### Tests file

The tests file contains information on the different descriptors of experimental conditions. *Exposure type* (`exposure_type`) refers to the method by which contact between the organism and the chemical is achieved. We only retain the exposure types that are most common and ensure consistent experimental designs^1^. *Media type* (`media_type`) is the liquid, in which the experiment was carried out. This feature was reduced to only contain fresh water and salt water experiments, also with the intention to remove tests conducted in environmental water samples that are basically non-reproducible and could have an effect by themselves.

The tests table contained other features that we retain but do not deem informative for modeling purposes. For *control type* (`control_type`), many of the data points are either not encoded or it is nontransparently reported what the encoding entails. The *test location* (`location`) was “laboratory” for more than 98% of the data points. *Application frequency unit* (`application_freq_unit`), describing how often the chemical was applied/renewed in the system, had such a high variability that it was not possible to normalize to a common unit. Most of the *organism life stage* (`organism_lifestage`) levels are uniformative.

##### Results file

The results file contains the information related to each result, *e.g.,* endpoint, effect, and concentration.

Mostly, *concentrations* (`conc1_mean`) are given as specific EC50 values. For limit tests or if the EC50 was outside the tested concentration range, the values are only given as above or below a concentration range (`conc1_mean_op` is either *<*, *≤*, *≥*, *>*). These values were removed. Other entries do not contain a mean effect value but a minimum and a maximum value (`conc1_mean_min` and `conc1_mean_max`, respectively). We averaged these two values if they were within one order of magnitude.

Effect concentrations are given in a variety of units (`conc1_mean_unit`). We only kept the units related to a mass or molar concentration and unified them to *mg/L*, from which we calculated the molar concentration in *mol/L*.

Similarly, *experimental durations* (`obs_duration_mean`) are given in a number of different units. We converted them to hours and retained 24, 48, 72, and 96 hours experimental periods to cover highly standardized short-term toxicity scenarios. Generally, during a 96 hour experiment effects are also recorded at the previous time points. So, often, later time points are an extension of previous ones, *i.e.,* repeated measurements in the same test. We checked several repeated tests and could not definitively determine that with similar experimental properties later measurements are consistently more or less sensitive than previous ones, even though this may be disguised in the general data variability. Thus, we included all such data points. If needed, one can filter to the latest data point for an experiment that is uniquely identified by the reference number.

For the *concentration type* (`conc1_type`), we retained all the entries with the levels “active ingredient” (A), “formulation” (F), and “total” (T). Chemicals, *i.e.,* the active ingredient that is of interest in toxicity testing, are often combined with other substances depending on their use case to delay or enhance their activity or facilitate transport to the target area/organ, *e.g.,* into a plant or adhering to a surface. We could not conclusively deduce that either of the concentration types is consistently more toxic than another and thus decided to include this information into our dataset.

##### Media characteristics file

The media characteristics file contained temperature (`media_temperature_mean`) and pH (`media_ph_mean`) values for some of the experiments. We removed or corrected unrealistic values (see next paragraph), calculated the average values if only minimum and maximum value were available, and converted all temperatures to °C.

##### Manual filtering of unrealistic values

The original dataset contains several values that required manual evaluation and cleaning/correcting. The manual filtering is documented in the code repository https://renkulab.io/gitlab/mltox/adore in the file *src/preprocessing.py* using the functions *remove_and_correct_entries*, *correct_ph_temp_entries* and *remove_invalid_test_entries*.

Some toxicity values were found to be either very high or very low, *i.e.,* EC50/LC50 *≥* 10^5^ *mg/L* and *≤* 10*^−^*^5^ *mg/L*. Also, some values deviated strongly from the other entries of the same experimental setting. We considered experimental settings with at least 25 repetitions and entries if their toxicity value was outside 3 times the interquartile range of the first and third quartile (based on the “outlier” definition of boxplots). These values were manually evaluated by checking against the original source. If the original source was not accessible, the values were checked against comparable data in the ECOTOX database, ideally from the same chemical and the same species. If that was not available, they were checked against the same chemical and a species from the same taxonomic group. If that was also not available or the values differed by several orders of magnitude, the data point in question was removed. Some data points were excluded because the toxicity values were obtained in multi-stressor scenarios, *e.g.,* testing chemical sensitivity in concert with salinity tolerance of a species or under non-representative conditions, *e.g.,* application of a chemical in orange juice as carrier medium^30^. We concluded that these toxicity values are not representative of the chemical’s toxic potential. We are aware that experimental setups like these likely make up part of the rest of the dataset as well and that they contribute to the high variability of effective concentrations (Figure 2), but manually checking all values is not feasible and would negate the use of an already assembled dataset, such as the ECOTOX database.

**Figure 2.**
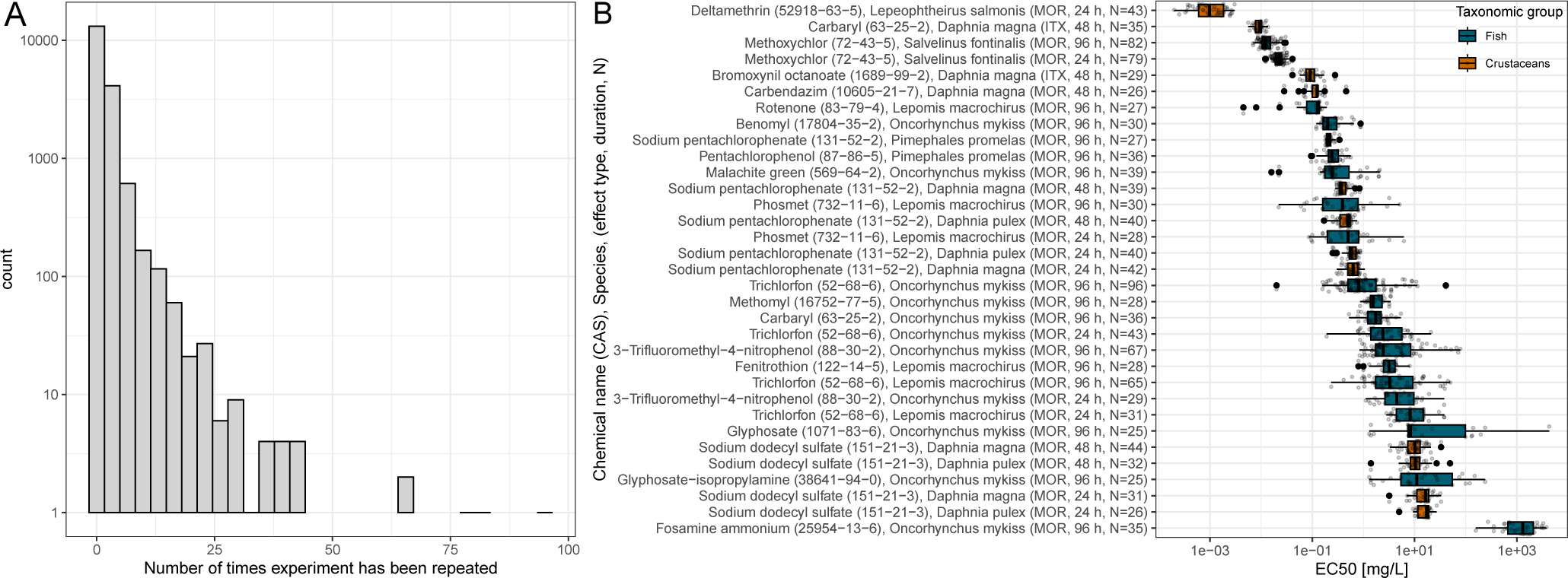
A: Histogram of the number of data points associated with a combination of chemical, species, and observation duration, concentration type, exposure type, media type, and effect type in our dataset. B: Boxplot for the distribution of toxicity values for experimental conditions with at least 25 values.

Similarly, we found unrealistic values for the experimental properties pH of the medium and temperature, *i.e.,* pH values smaller than 4 and larger than 10, and temperatures exceeding 50 °C. Again, we checked the values manually against the original literature and excluded cases where (waste)water samples with extreme pH were the test medium and where the pH was strongly influenced by the test substance. For temperature, we employed expert judgment and concluded that the values were mostly conversion errors, *e.g.,* unit given as °C, when actually °F, or decimal errors, *e.g.,* 250 °C when actually 25.0 °C, and adjusted the values accordingly.

Theoretically, it should be impossible to derive an EC50 value larger than the chemical’s water solubility. Nonetheless, a number of data points contain EC50 values that lie above the given water solubility of the compound. This can partly be explained by most water solubility values being based on predictions or derived from experiments based on ultrapure water, while exposure media already contain several micronutrients and salts. One option on how to deal with these entries is to remove all data points for which the effective concentration is higher than the chemical’s water solubility^31^. Alternatively, as was done for the EnviroTox database, entries where the toxicity value exceeds the water solubility by a factor of 5 are specifically marked, but not removed^16^. The EC50 values in our dataset are calculated from the nonlinear regression on the concentration-response-curves and therefore themselves come with inherent uncertainty. Accordingly, all data points are uncertain to one degree or another, which in this case only becomes apparent for the data points where the EC50 exceeds water solubility. Therefore, we decided against applying this filtering step.

### Species-related data

We include two kinds of species-related data, species-specific ecological, life-history, and “pseudo“-data related to the DEB modeling framework and phylogenetic distances between species.

#### Add my Pet

Species-specific data was acquired from the Add my Pet collection (AmP), a manually curated database linked to the dynamic energy budget (DEB) theory framework^32^, that contains information for more than 3,400 animals (August 2022). The DEB theory is based on heterotrophs, *e.g.,* animals like fish and crustaceans, and does not cover autotrophs such as algae. Therefore, comparable data on algae is not included in our dataset. We downloaded the AmP collection (dated November 25^th^, 2022) and extracted information for the species present in the core dataset. This includes three types of information: Ecological data, life history data, and pseudo-data. The included features are shown in Table 2.

**Table 2.** Add my Pet features and their encoding. See Annex Figure 12 for more detailed descriptions of these features.

| type | feature | code | unit |
| --- | --- | --- | --- |
| ecology | ecozone |  |  |
|  | climate |  |  |
|  | migration |  |  |
|  | food |  |  |
| life history | life span | am | *d* |
|  | body length at birth | Lb | *cm* |
|  | body length at puberty | Lp | *cm* |
|  | ultimate body length | Li | *cm* |
|  | reproductive rate | Ri | *#/d* |
| pseudo-data | allocation fraction to soma | kap | - |
|  | energy conductance | v | *cm/d* |
|  | volume-specific somatic maintenance cost | p_M | *J/d · cm<sup>3</sup>* |

##### Ecology

This data category includes information about ecological peculiarities and lifestyle of a species. We selected climate zone and ecozone, *i.e.,* the climatic and geographical area, respectively, where the species occurs (Annex Table 8 and Annex Table 9). Additionally, we selected migratory behavior, *e.g.,* anadromous/catadromous species that migrate up-/downstream for reproduction (Annex Table 10), and food type, *e.g.,* carnivorous, herbivorous, parasitic, *etc.* (Annex Table 11).

For three features, we reduced the complexity of the original encoding. The AmP collection provides 51 different levels for climate zone and 56 for ecozone, respectively. Similarly, the food encoding has 77 different levels. These features follow a hierarchical encoding, which was reduced by retaining only the top level. Further information on the encodings can be found in the Annex Tables 8-11 and on the AmP website.

##### Life history

The AmP collection also provides life history data, which are usually taken from literature, but sometimes estimated or extrapolated from related species. As a consequence, some features are not available for each species and the resulting dataset contains missing values. We included the features which had the highest coverage of number of data points with the core dataset: life span, body length at birth, body length at puberty, ultimate body length, and maximum reproductive rate. Life span and ultimate body length are available for all species.

##### Pseudo**-data**

Pseudo-data are species-specific parameters related to the modeling framework that the AmP collection forms the basis of. We included the three pseudo-data features showing reasonable variability across species: allocation fraction to soma^33^, energy conductance, and volume-specific somatic maintenance cost. These features are available for every species included in the AmP database. Definitions are given on the DEB-portal and in Kooijman *et al.* (2008)^34^.

#### Phylogenetic distances from TimeTree

Since closely related species often are similarly sensitive to the same chemicals compared to distantly-related species^35^, we include the phylogenetic distance between species as possible modeling features. The phylogenetic distance represents how closely related a pair of species is to each other by the time since they diverged from the last common ancestor^36^. We used TimeTree Version 5 to produce a phylogenetic tree of the species in our data. The tree was then transformed into a phylogenetic distance matrix using the Python package `dendropy`^37^.

##### Chemical data

We include several chemical features of which some are intended as modeling features, namely chemical properties and molecular representations. The others should only be used to help interpret modeling results, namely the ClassyFire ontology and the functional use categories.

#### Chemical properties

We added basic chemical properties from DSSTox^38^, the PubChem database^28^ and through the Python package RDKit^39^. The properties are molecular weight (MW in *g/mol*), water solubility (WS in *mg/L*), melting point (MP in °C), number of heavy atoms (*i.e.,* all atoms excluding hydrogen), number of bonds, double bonds, and triple bonds, number of ring structures, and the number of OH groups. Additionally, we included the octanol-water partition coefficient (logP, sometimes referred to as *logK_ow_*, the logarithm of the relative solubility of a compound in the solvents octanol and water^40^, positive/higher values, indicate a higher lipophilicity of a compound), and the acid dissociation constant (pKa).

The pKa regulates at which pH a substance dissociates. Since, for some compounds, the pH of the medium influences the toxicity^41^, it is reasonable to include the pKa as a feature to be used together with the medium’s pH. The pKa values were obtained through the OECD QSAR toolbox (V4.5)^42^ via the SMILES. When several predictions were available, we used their median. Median values above 30 were excluded.

#### Molecular representations

Presenting chemical structures to a ML model such that it can be used in an informative manner poses a challenge. Standard molecular formats such as connection tables and SMILES are not valid input for most ML models. Chemicals need to be provided in a form “understandable by an algorithm” which Staker *et al.*^43^ refers to as a molecular representation. We used three well-established molecular fingerprints^44^: MACCS^45^, PubChem^46^ and Morgan^47^ fingerprints, a fingerprint developed for toxicity predictions, ToxPrint^48^, the molecular descriptor Mordred^49^ as well as the mol2vec embedding^50^.

Molecular representations can be divided into non-learned and learned representations^43^. Besides computable features *i.e.,* chemical properties, which we introduced in the previous section, molecular fingerprints are widely used non-learned representations. Fingerprints are fixed-length, binary vectors where each bit indicates presence (“1”) or absence (“0”) of a substructure. MACCS, consisting of 166 publicly available bits, PubChem, consisting of 881 bits, and ToxPrint, consisting of 729 bits, are substructure keys-based, *i.e.,* each bit corresponds to a predefined substructure^2^. The Morgan fingerprint is a circular fingerprint based on all fragments that can be extracted within a bond radius around each atom of the molecule. These fragments are then hashed to create fingerprints comparable across chemicals of different size. We used RDKit^39^ to calculate the MACCS and Morgan fingerprints for all chemicals in the dataset. The Morgan fingerprint was calculated with a radius of 2 bonds and hashed to 2048 bits. The PubChem fingerprint was extracted as CACTVS fingerprint using PubChemPy^29^.

The ToxPrint was extracted using ChemoTyper editor v1.1 and ToxPrints v2.0^48^. Mordred is a collection of molecular descriptors including atom and bond counts, chemical properties like molecular weight and logP, and properties calculated from the 2D and 3D structures^49^. These descriptors are calculated using the Python package mordredcommunity^51^, a community effort to continue the initial software implementation. We included the 719 molecular descriptors which could be calculated for all chemicals in the preprocessed dataset.

The last representation is learned. Jaeger *et al.* developed the molecular embedding mol2vec, which applies the word embedding word2vec^52^ on substructures derived from the Morgan algorithm. This embedding treats substructures as if they were words, and molecules as if they were sentences^50^. Jaeger *et al.* trained a 300 dimensional embedding on 19.9 million compounds, which is available as a Python package via https://github.com/samoturk/mol2vec and was used to calculate 300-dimensional feature vectors for all chemicals in the dataset.

#### Chemical ontology from ClassyFire

Chemical structures can be organized into a taxonomical system through the standardized classification system ClassyFire^53^. This information can either be retrieved using the ClassyFire API or the Python package pybatchclassyfire based on InChIkeys. We consider this feature more useful for the interpretation of modeling results than for the modeling itself.

#### Functional use categories

The US EPA provides a Chemicals and Products Database (CPDat) containing product applications and functional uses for various chemical compounds (December 2020 Release). The database provides functional uses in two variants. The OECD function is an internationally harmonized functional use category^54^, which is suitable for most chemicals in our dataset except for the category “biocide”. Biocides can be subdivided in insecticides, herbicides, fungicides, antimicrobial agents, and others, which is relevant for our dataset. This information is contained in the reported functional uses, which are unharmonized data entries. Please refer to the official documentation^54^ for definitions of the OECD functions.

We processed the data by combining closely related OECD functions as indicated by the description^54^, *e.g.,* dye and pigment are renamed to colorant. The reported functional uses are harmonized in several steps: All entries are set to lower case and unnecessary white space is removed. Non-informative entries such as “not reported”, “proprietary”, or “impurity” are removed. The other entries are unified where possible, *e.g.,* entries starting with “herbicide” are renamed to only “herbicide”, or entries containing the term “catalyst” are renamed to “chemical reaction regulator” as is the name of the corresponding OECD function. Then, reported functional uses longer than 30 characters or occurring less than four times are removed. This results in 43 OECD functions and 174 reported functional uses. For most chemicals, there are several OECD functions and reported functional uses. We provide all of them as well as only the most prevalent ones, *i.e.,* the uses making up more than 20% of the entries for a compound.

The functional use categories should *not* be included in modeling, but only be used to enable a better understanding of modeling results. In fact, its inclusion would constitute a form of data leakage, since, *e.g.,* labeling a substance group as biocides would correlate well with higher toxicity towards certain species, but the information was not transmitted through a chemical feature but through this label.

### Further data processing

#### Overlap between data categories

The data categories, described in the previous sections, did not necessarily contain a feature value for every data point, *e.g.,* the ECOTOX data contains 1,690 species and 4,772 chemicals after processing but only data for 665 and 270 of those species are available in TimeTree and AmP, respectively.

This leads to a trade-off between retaining more data points and expanding the feature space. The longest version of the dataset contains more data points but has fewer features, whereas the dataset with the highest number of features retains a far smaller number of data points. Since the bottleneck for environmental risk assessment models is usually the scarcity of data, we prefer a bigger dataset to a larger feature space and therefore provide a long core dataset.

Apart from using ML to predict toxicity, we envision our dataset being used for the exploration of other ecotoxicologically relevant research questions. Therefore, we provide incomplete data tables, *i.e.,* containing missing values for some features, which are intended for future studies that are not tied to the challenges proposed and discussed in this paper.

#### Filter by taxonomic group and effect group

After all input data was harmonized and pre-filtered, and taxonomic information as well as chemical properties were added, we filtered the data to only contain acute mortality experiments for the three taxonomic groups fish, crustaceans and algae. For each taxonomic group, this encompasses the corresponding effect groups (Table 1). The endpoints were filtered to only include LC50 and EC50.

#### Final filtering

The processed data on acute mortality was then filtered to obtain a dataset suitable for modeling. To employ the mol2vec embedding and compare it to the other molecular representations, we only keep compounds which are compatible with it^50^. Also, we only keep compounds for which water solubility is available as we deem it a potentially important modeling feature. Additionally, we remove data points without phylogenetic and AmP information. These filtering steps remove around 50% of the processed data and leaves us with a dataset comprised of predominantly organic chemicals.

##### Data splittings

Data splitting, *i.e.,* the generation of training and test data subsets, and of cross-validations folds, can greatly affect model performance. Possible causes are the inherent variability of the data itself and (non-obvious) data leakage. The latter describes a case where information about the target is introduced which should not legitimately be used for modeling^55^. Data leakage has also been described as a “spurious relationship between the independent variables and the target variable”^10^ and leads to overestimated model performance^55^.

We discuss different data splitting schemes, of which most account for some sources of data leakage. Ideally, a combination of chemical and species would only appear in either the training or the test set. Since our dataset is skewed both with respect to chemicals and species, splits by chemical *and* species would lead to highly imbalanced subsets and are not feasible. We can only reasonably apply a split by either chemical or species, *i.e.,* all data points belonging to a chemical or species, respectively, are found in either the training or the test set. We focus on splits by chemical, due to the higher variability in chemicals compared to species and because extrapolating across chemicals allows us to address the problem of predicting toxicity of untested or yet unknown chemicals.

##### Split totally at random

The simplest train-test-split can be achieved by random sampling of data points, provided by us as *totally random*, which has been the main approach in previous work applying ML to ecotoxicology, and generally suffices for a well-balanced dataset without repeated experiments^31,56,57^. For our dataset with repeated experiments, *i.e.,* data points coinciding in chemical, species, and experimental variables (Figure 2), the *totally random* approach has a high risk of data leakage and the associated overestimated model performances. With this splitting, the same chemical is likely to appear in both training and test set.

Thus, we present stratified splits as alternatives which ensure that chemicals are not shared between training and test set, and not between cross-validation folds. We provide splits for 5-fold cross-validation.

##### Splits by chemical compound

The straightforward approach is to split by chemical compound, ensuring that a chemical is put into either the training or the test set. We provide two variants here, where compounds are split at *random by chemical compound* or *by occurrence of chemical compounds*. For the latter, compounds are sorted by the number of experiments performed on them, *i.e.,* those with most experiments at the top. Then, the first five compounds are put into the training set and the sixth is put into the test set. This is repeated with the subsequent compounds until all are distributed. The five cross-validation folds are filled accordingly, *e.g.,* the most common compound goes to fold 1, the second most common to fold 2, and so on. With both variants, it is likely that similar chemicals are shared between the training and test set, and between the cross-validation folds.

##### Splits by molecular scaffold

Molecular scaffolds can be used to categorize molecules based on their core structure, *i.e.,* the molecular backbone, notwithstanding side chains and functional groups. Scaffold splitting has been used in drug discovery^58^ and was deemed to lead to more optimistic modeling outcomes than relying on random splitting^59^. We utilize two scaffold types: the Murcko scaffold and the generic scaffold. The Murcko scaffold reduces a molecular structure to ring systems and the structures connecting these ring systems^60^. The generic scaffold goes one step further by replacing all heteroatoms in the Murcko scaffold with carbon atoms and all bonds with single bonds. Both scaffold types were calculated using RDKit^39^.

For both scaffold types, we provide several data splits. Scaffolds are put at random into either training or test set (*scaffold-murcko* and *scaffold-generic*). Here, we follow the procedure described by Yang *et al.* and put molecules whose scaffold is more common than half the test size into the training set^58^. The others are distributed at random between training and test set. Additionally, we provide two leave-one-out (LOO) splits where we put either the most common scaffold (*scaffold-murcko-loo-0* and *scaffold-generic-loo-0*) or the second most common scaffold (*scaffold-murcko-loo-1* and *scaffold-generic-loo-1*) in the test set and the others in the training set. Also, we include a “leave-last-out” (LLO) alternative, where we put the least common scaffolds in the test set, until the test size is reached, and the remaining ones in the training set (*scaffold-murcko-llo* and *scaffold-generic-llo*). The last splitting scheme represents a generalization to new compounds with more exotic structures. We provide several splits by both scaffold types since this is an area of active research and it is not yet known when and if splits by scaffold should be used.

We provide 11 splits, including cross-validation folds if applicable, for the data subsets where we train and test on the same taxa. For the data subsets where we train on crustaceans and algae and test on fish, we do not provide splits by molecular scaffold. Cross-validation is applicable for *totally random*, *random by chemical compound*, *by occurrence of chemical compounds*, *scaffold-murcko* and *scaffold-generic* but not for the LOO and LLO options. Since the split *by occurrence of chemical compounds* puts one sixth, *i.e.,* 17%, of the data points in the test set, we maintain the associated ratio of 83:17 for all train-test-splits to have comparable sizes across the data subsets.

## Data Records

The dataset is available on the Eawag Research Data Institutional Repository (ERIC), the institutional data repository of Eawag under https://doi.org/10.25678/0008C9. In the final dataset, there are three folders (*processed*, *chemicals*, *taxonomy*). All files are provided as csv or tsv files. All data files, including the raw files, are also part of the code repository https://renkulab.io/gitlab/mltox/adore.

Intermediate files as well as the final files, one for each prediction challenge, can be found in the folder *processed*. We include the intermediate files such that the user can follow the processing steps. They encompass the processed separate files from ECOTOX for species (*ecotox_species.csv*), tests (*ecotox_tests.csv*), results (*ecotox_results.csv*), as well as compiled chemical properties (*ecotox_properties.csv*) in the subfolder *before_aggregation*. These files are then combined to get *ecotox_mortality_processed.csv* and then filtered to get *ecotox_mortality_filtered.csv*.

For each challenge, we provide a data subset which starts with the name of the challenge as given in Table 3 and ends with *_mortality.csv*. These datasets contain all information needed for modeling and further columns to uniquely identify and retrace each entry (Annex Table 6).

**Table 3.** Prediction challenges. The names of the challenges on the whole dataset start with “a”, those within a taxonomic group start with “t”, and those on a single species start with “s”. The challenge “a-CA2F-diff” generalizes both to species and chemicals whereas “a-CA2F-same” only generalizes to species. In contrast, “a-FCA2FCA” generalizes to neither of them.

| Name | Train on | Test on | Data split |
| --- | --- | --- | --- |
| a-CA2F-same | crustaceans and algae | fish | by occurrence of chemical compounds |
| a-CA2F-diff | crustaceans and algae | fish | by occurrence of chemical compounds |
| a-FCA2FCA | all data | all data | by occurrence of chemical compounds |
| t-F2F | fish | fish | by occurrence of chemical compounds |
| t-C2C | crustaceans | crustaceans | by occurrence of chemical compounds |
| t-A2A | algae | algae | by occurrence of chemical compounds |
| s-F2F-1 | *Oncorhynchus mykiss* (Rainbow Trout) | *Oncorhynchus mykiss* | by occurrence of chemical compounds |
| s-F2F-2 | *Pimephales promelas* (Fathead minnow) | *Pimephales promelas* | by occurrence of chemical compounds |
| s-F2F-3 | *Lepomis macrochirus* (Bluegill) | *Lepomis macrochirus* | by occurrence of chemical compounds |
| s-C2C | *Daphnia magna* (Water Flea) | *Daphnia magna* | by occurrence of chemical compounds |
| s-A2C | *Chlorella vulgaris* (Microalga) | *Chlorella vulgaris* | by occurrence of chemical compounds |

Additionally, we provide five separate files in the folder *chemicals*. The chemical ontology (*classyfire_output.csv*) and the functional use categories (*functional_uses_output.csv*) can be matched with the data in the challenges files using the InChIkey or the DTXSID, respectively. Three files are explaining the bits of MACCS, PubChem and ToxPrint (*maccs_bits.tsv*, *pubchem_bits.csv*, *toxprint_bits.csv*). The file *tax_pdm_species.csv* in the folder *taxonomy* contains phylogenetic distance information which can be used for modeling.

## Technical Validation

Here, we describe the dataset and its features. Information on how the dataset was compiled can be found in the “Methods” section. We provide several data subsets related to the proposed prediction challenges presented in the “Usage Notes” section.

The figures and numbers given in this section refer to the comprehensive file *a-FCA2FCA_mortality.csv*.

### Ecotoxicological Data

#### Toxicity

The outcome of ecotoxicological experiments is a continuous value, in our case an EC50 value, or LC50 for mortality. The distributions of the different effects for the three taxonomic groups are shown in Figure 3A. The overall spread of EC50 values is comparable across effect groups, especially for fish and crustaceans. Physiology, only recorded in algae, has a bias towards low EC50 values, which is reflected in Figure 3C, where the distribution across toxicity classes is comparable to that across taxonomic groups (Table 4), while few compounds were non-toxic to algae. This could entail a bias in the compounds tested on algae, either with the expectation of high toxicity or because algae are more sensitive than the other taxonomic groups. Similarly, since the database has grown over time, it could hint to historical shifts related to the inclusion of algae as model organisms. Annex Figure 10 confirms that the first reported tests on algae was done in 1974, 28 years and 10 years later than for fish and crustaceans, respectively. Figures 3B and 3C visualize the distribution of data points across the three taxonomic groups for either a binary or a multi-class classification, respectively.

**Figure 3.**
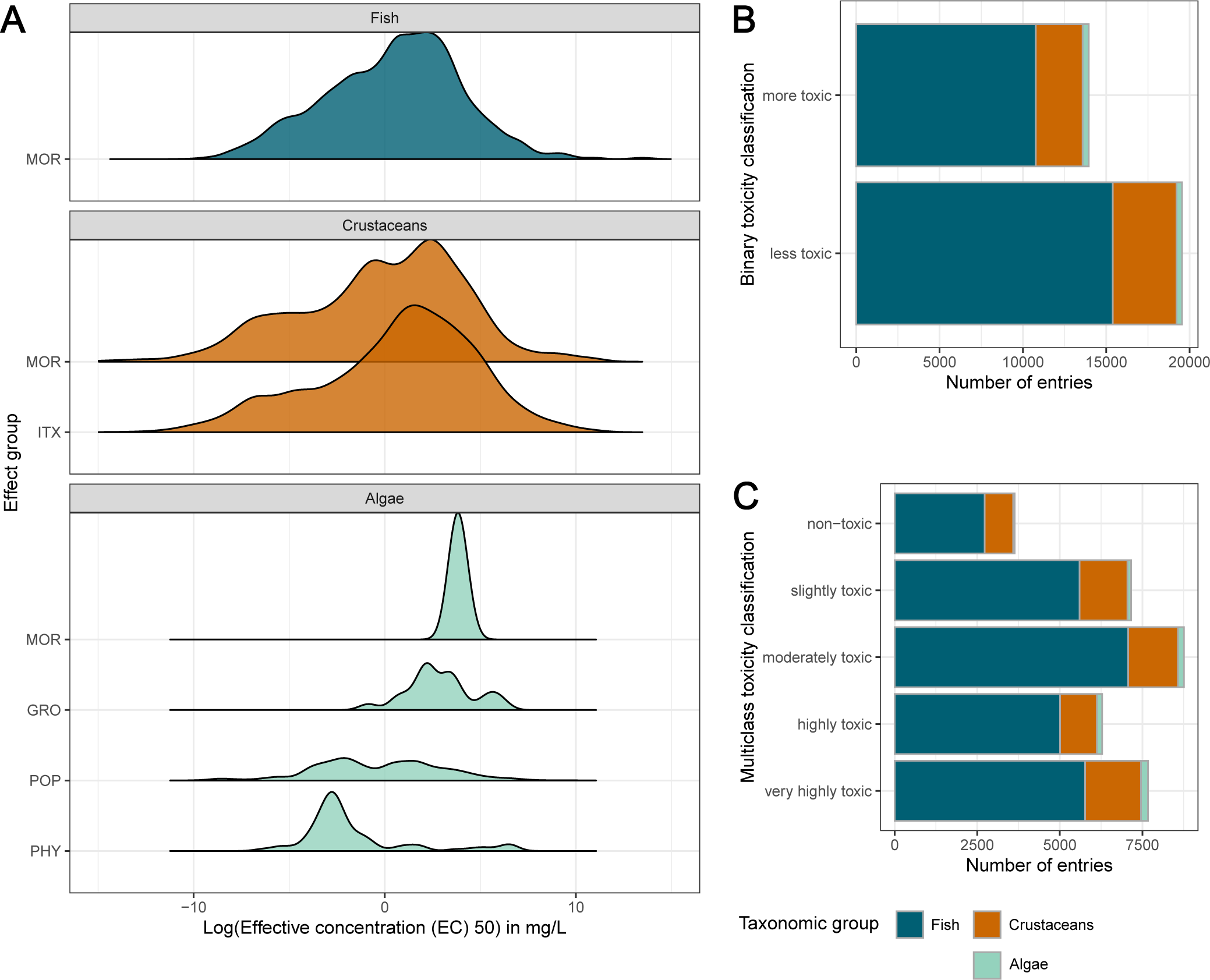
A: Overview of the toxicity distributions across effect groups by taxonomic groups. B, C: Toxicity classes across taxonomic groups in a binary and 5-level classification system, respectively.

**Table 4.** Total number of data points after processing. N: number of data points.

| Taxonomic group | Number of species | N | % |
| --- | --- | --- | --- |
| Fish | 140 | 26,114 | 78.1 |
| Crustaceans | 17 | 6,630 | 19.8 |
| Algae | 46 | 704 | 2.1 |
| **All** | **203** | **33,448** | **100.0** |

#### Repeated experiments and reproducibility

As we are dealing with highly variable *in vivo* data, experiments with the same experimental conditions can lead to differing results. The dataset contains a large fraction of “repeated experiments”, *i.e.,* data points that overlap in species, chemical, and the experimental parameters effect type, media type, observation duration, exposure type and concentration type. As a consequence, our modeling target EC50 rather resembles a probability distribution than a singular ground truth. We show this in Figure 2A with a histogram of the number of data points sharing a parameter combination. Most experiments have only been done once or repeated a few times. There are three experimental settings that have been repeated around 80 times each: the pesticide Trichlorfon (CAS number 52-68-6) has been tested on the rainbow trout (*Oncorhynchus mykiss*) 96 times while the insecticide Methoxychlor (72-43-5) has been tested on the brook trout (*Salvelinus fontinalis*) over an experimental duration of 24 and 96 hours, respectively, 79 and 82 times. Some of the experimental outcomes vary up to several orders of magnitude, even within this restrictive set of parameter values, as can be seen in Figure 2B, which shows experiments repeated at least 25 times. This variability needs to be considered when evaluating model predictions originating from this data.

#### Experimental features

Experimental features like media type and observation duration may influence the experimental outcome. See Annex Figure 11 for an overview of the features and their levels.

## Species-related data

In total, there are 203 species across all taxonomic groups of which fish make up around two third (Table 4). A few species have been tested much more often than most, causing the dataset to be skewed with respect to species. For fish, those are rainbow trout (*Oncorhynchus mykiss*), fathead minnow (*Pimephales promelas*), and the bluegill sunfish (*Lepomis macrochirus*) (Figure 4A). By far the most tested crustacean species is the waterflea *Daphnia magna*, followed by another water flea species, *Daphnia pulex* (Figure 4B). For algae, the microalgae *Chlorella vulgaris* and the green algae *Chlamydomonas reinhardtii* are the most abundant species in the dataset (Figure 4C).

**Figure 4.**
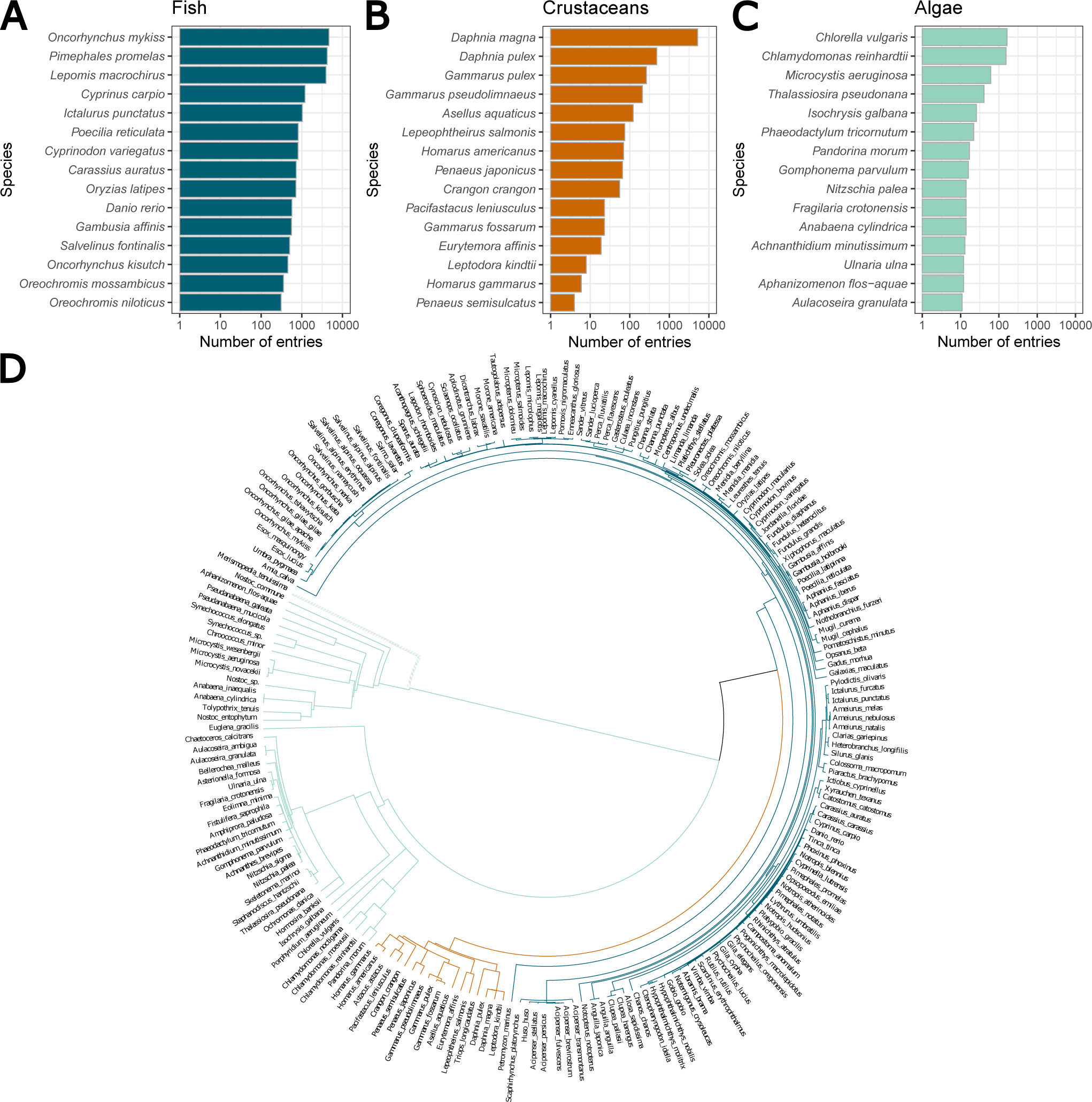
A-C: Overview of the top 15 species per taxonomic group. D: The phylogenetic distances as circular dendrogram. Line lengths after branching indicate phylogenetic distances.

Generally, apart from the dominance of a number of model species, the diversity of organisms included in the dataset is high and many taxonomic sublevels are present. A dendrogram of the species visualizing the phylogenetic distances between species are shown in Figure 4D. The length of lines after branching indicates the distance between species. This is most striking in the early branching of the algae from the two animalia groups fish and crustaceans (black line). Even though we consider it a good indicator of species relatedness, the user needs to consider that the phylogenetic distance matrix cannot straightforwardly be used with every type of model.

Other species-related data are focused on ecology, life-history and pseudo-data from the DEB theory framework, originating from the AmP database. The distributions of these species-specific features are shown in Annex Figure 12. They are only available for fish and crustaceans. Many species are found in several climate and ecozones, or accept different kinds of food. Therefore, we included all these combinations of levels in the data. The level abbreviations are divided by an underscore and are meant to be reformatted into multi-hot-encoded matrices for modeling use (Annex Figure 12A). Definitions for the encoding levels are provided in the Annex Tables 8-11. The life history data and the pseudo-data are given in Annex Figure 12B. The histograms are dominated by the variability in fish species whereas crustaceans cover smaller ranges. This is more prominent for the life history features, body length at three stages and reproductive rate, than for the pseudo-data features. The dataset contains many fish species of which some grow large, whereas the covered crustaceans all belong to fairly small-bodied taxonomic orders.

The importance of these parameters for modeling is difficult to foresee but we are positive that information distinguishing species is valuable to at least partly explain variation in the response of animals.

## Chemical data

In total, the dataset encompasses ecotoxicological tests on 2,408 chemicals. Substances are routinely tested on several species and taxonomic groups, which is reflected in the Venn diagram in Figure 5A. 111 chemicals have been tested in all three taxonomic groups, while 1,905 and 1,444 have been tested on fish, and crustaceans, respectively, while only 174 chemicals have been tested on algae. Likewise, Figure 5B visualizes the imbalance between the number of chemicals tested on different pairings of taxonomic groups and the number of data points for these tests. As an example, the 111 chemicals tested in all taxonomic groups represent 10,568 data points, while the 452 chemicals tested just on crustaceans only make up 825 data points.

**Figure 5.**
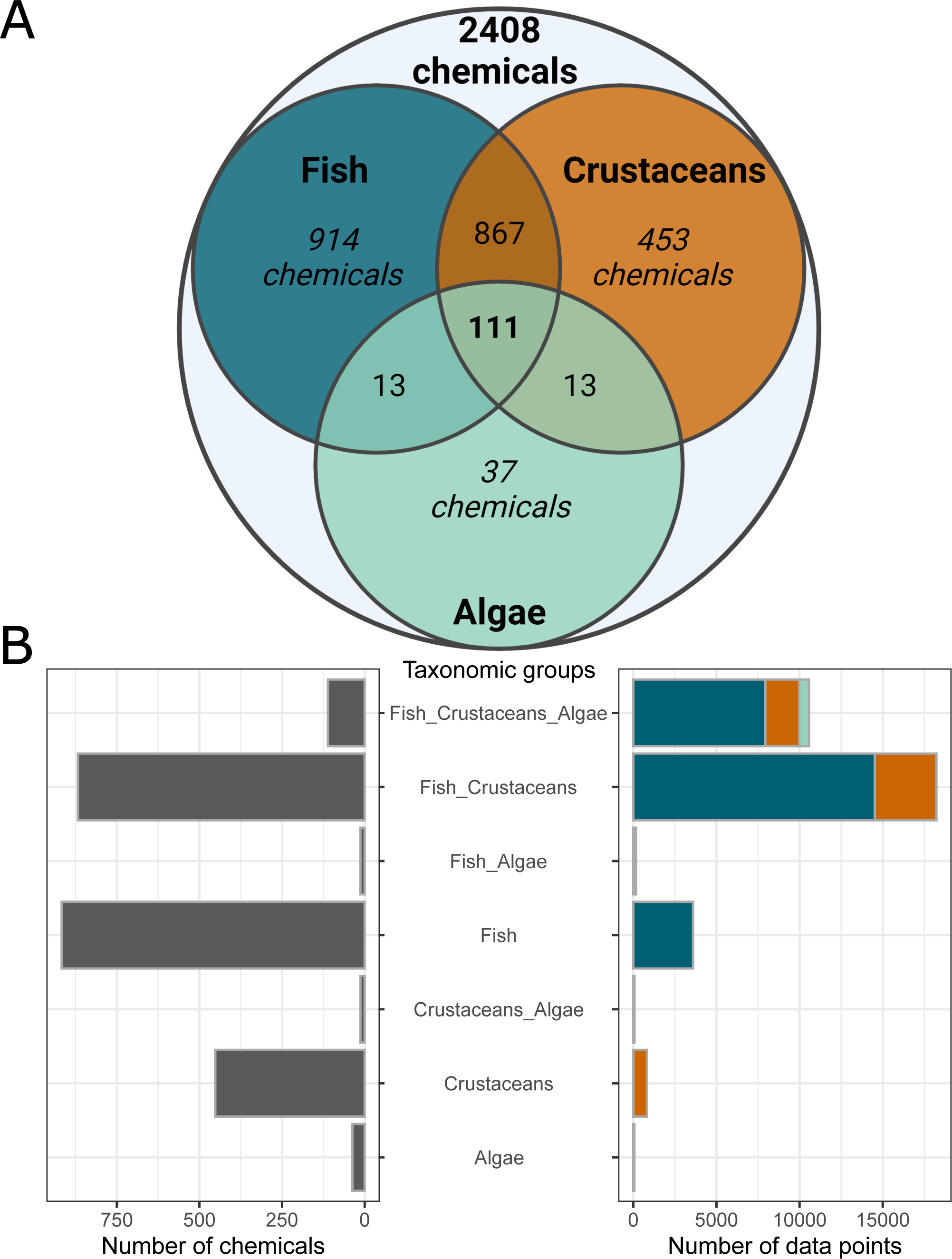
A: Venn diagram of the 2,408 tested chemicals. As reading example, 914 chemicals were tested only on fish, 867 chemicals were tested on fish and crustaceans, and 111 chemicals were tested on all taxonomic groups. B: Overview of the number of chemicals tested (left) in the different combinations of taxonomic groups and the related number of data p**_2_**o**_1_**i**_/_**n**^4^**t**^0^**s (right), indicating that many tested chemicals do not automatically translate to many data points.

As with species, some chemicals have been tested much more often than others. Figures 6A-C show the number of data points by taxonomic group for the top 15 chemicals with the most associated data points. Primarily, chemicals are tested on fish, then on crustaceans, and lastly on algae. The insecticide Carbaryl (63-25-2) has been tested most often on fish with almost 600 data points, whereas the pesticide Sodium Pentachlorophenate (131-52-2) has been tested most often on crustaceans. For algae, the herbizide Atrazine (1912-24-9) makes up around 15% of the data points.

**Figure 6.**
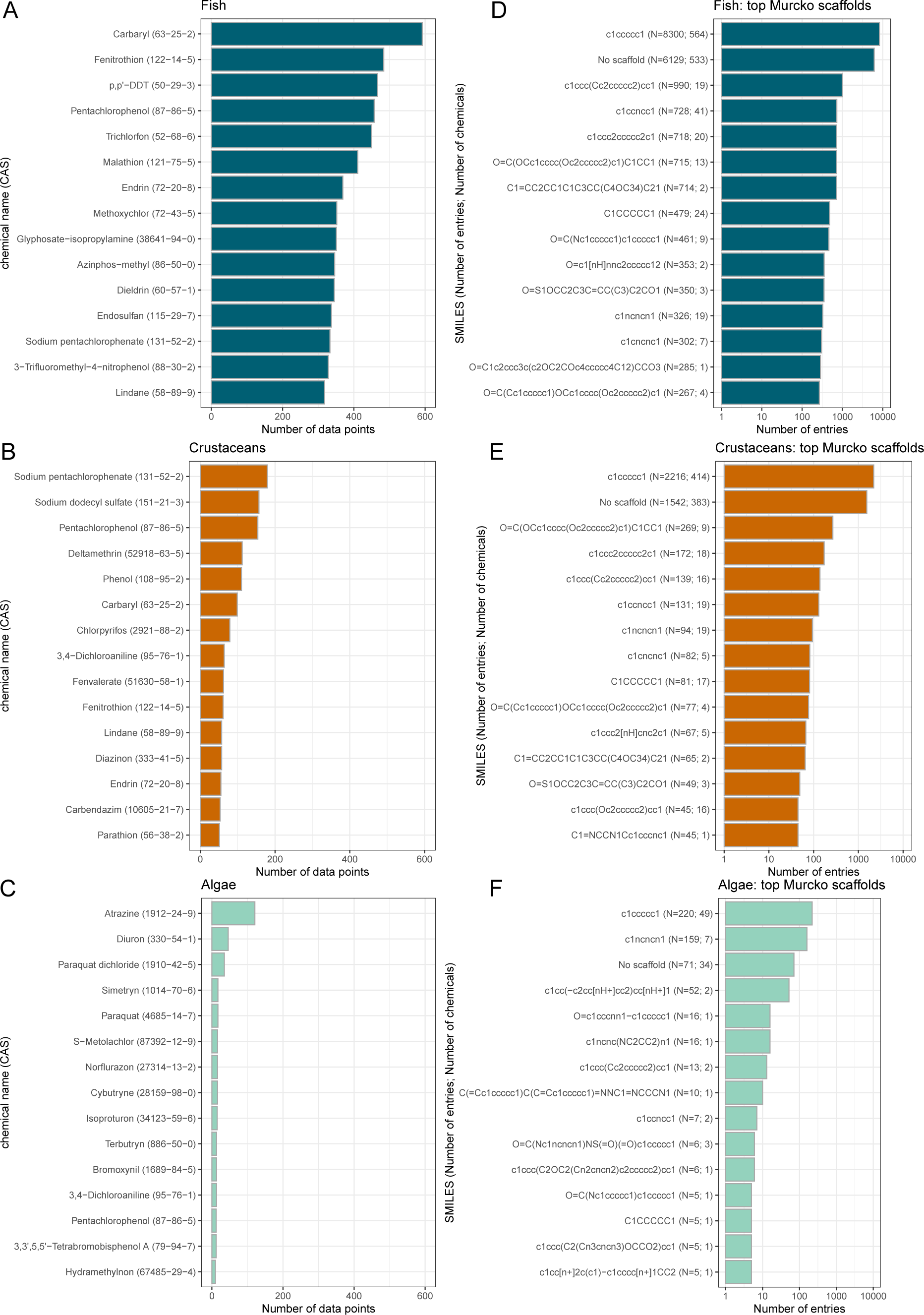
Top chemicals (A-C) and Murcko scaffolds (D-F) for the three taxonomic groups. Annex Table 12 gives an overview of the structures of each scaffold.

To better understand the chemical space of our dataset, we look at pairwise similarities between chemicals. The Tanimoto similarity was shown to be an appropriate choice for similarities based on molecular fingerprints^61^ and is given as a value between 0 (dissimmilar) and 1 (equal). We calculated the similarity using the MACCS fingerprint with the BulkTanimotoSimilarity function from the Python package RDKit. Annex Figure 13 depicts a histogram of all pairwise comparisons of the 2,408 chemicals in our dataset, which shows that the majority of chemicals are fairly dissimilar, with a mean similarity of 0.085 as indicated by the dotted vertical line.

Understanding the chemical space can also be approached using the chemical ontology ClassyFire^53^, whose hierarchical structure for our dataset is shown in the icicle chart in Annex Figure 14. 2,250 of the 2,408 chemicals in the dataset could be categorized with ClassyFire. Our dataset mainly consists of organic compounds, which is a consequence of the final filtering step which only includes water soluble and mol2vec compatible compounds. The largest superclass of the 17 superclasses, benzenoids, in total makes up more than a third of the chemicals, followed by organoheterocyclic compounds, and organic acids and derivatives. The chemicals are further divided into 140 classes and 251 subclasses, thus indicating a wide variety of compounds in the dataset. The selected chemical properties are described in Table 5 and histograms shown in Annex Figure 15.

**Table 5.**
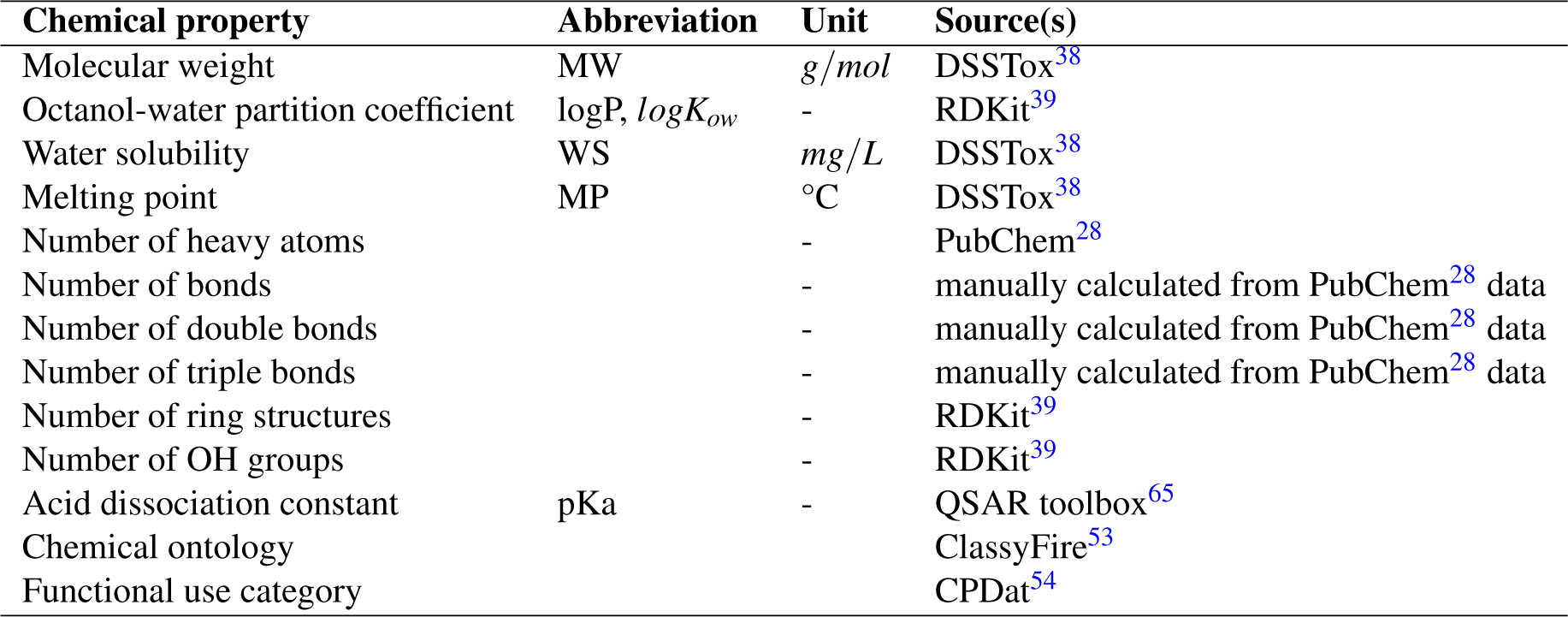
Overview of the chemical properties and their sources. CPDat: Chemicals and Product Database.

| Chemical property | Abbreviation | Unit | Source(s) |
| --- | --- | --- | --- |
| Molecular weight | MW | $g/mol$ | DSSTox<sup>38</sup> |
| Octanol-water partition coefficient | logP, $logK_{ow}$ | - | RDKit<sup>39</sup> |
| Water solubility | WS | $mg/L$ | DSSTox<sup>38</sup> |
| Melting point | MP | $^{\circ}C$ | DSSTox<sup>38</sup> |
| Number of heavy atoms |  | - | PubChem<sup>28</sup> |
| Number of bonds |  | - | manually calculated from PubChem<sup>28</sup> data |
| Number of double bonds |  | - | manually calculated from PubChem<sup>28</sup> data |
| Number of triple bonds |  | - | manually calculated from PubChem<sup>28</sup> data |
| Number of ring structures |  | - | RDKit<sup>39</sup> |
| Number of OH groups |  | - | RDKit<sup>39</sup> |
| Acid dissociation constant | pKa | - | QSAR toolbox<sup>65</sup> |
| Chemical ontology |  |  | ClassyFire<sup>53</sup> |
| Functional use category |  |  | CPDat<sup>54</sup> |

The ranges of the chemical properties indicated in the histograms are a simple approach to describe the applicability domain of models trained on the dataset. Predictions for chemicals whose properties fall outside of any of these ranges will not be trustworthy. More advanced approaches to characterizing the applicability domain are discussed elsewhere^62^.

Benzenoids being the most prevalent chemical superclass coincides with the findings from the Murcko scaffolds. There are 525 different Murcko scaffolds in our dataset of which a benzene ring and “no scaffold”, *i.e.,* the molecule does not contain a ring system, are by far most common for all taxonomic groups (Figures 6D-F). For algae, 1,3,5-Triazine (*c1ncncn1*) is more common than “no scaffold”. The latter is a very heterogenous group, since the only commonality among them is the lack of rings in their structure. The generic scaffold can be considered more reductionist by creating even broader scaffold categories. Here, the most common scaffolds are “no scaffold”, and cyclohexane.

Chemicals associated with different scaffolds produce different toxicity ranges which can be seen by the different toxicity ranges covered by the top 15 Murcko scaffolds (Figure 8). The structures of these 15 Murcko scaffolds are depicted in Annex Table 12. The two main scaffolds cover a fairly similar toxicity range, whereas less-represented scaffolds either mostly represent a narrower or sometimes a broader part of the toxicity range. Some scaffold are particularly toxic or non-toxic for certain taxonomic groups, *e.g.,* the *O=C(OCc1cccc(Oc2ccccc2)c1)C1CC1* scaffold is less toxic to algae than to fish and crustaceans. It is the backbone of pyrethroid insecticides, most notably Permethrin, Deltamethrin, and Cypermethrin. On the other hand, chemicals based on the 1,3,5-Triazine scaffold are more toxic to algae and include primarily herbicides such as Atrazine, Simazine, and Terbutryn. These findings, apparent from the raw data, underline the informational value contained in the grouping according to scaffolds.

**Figure 7.**
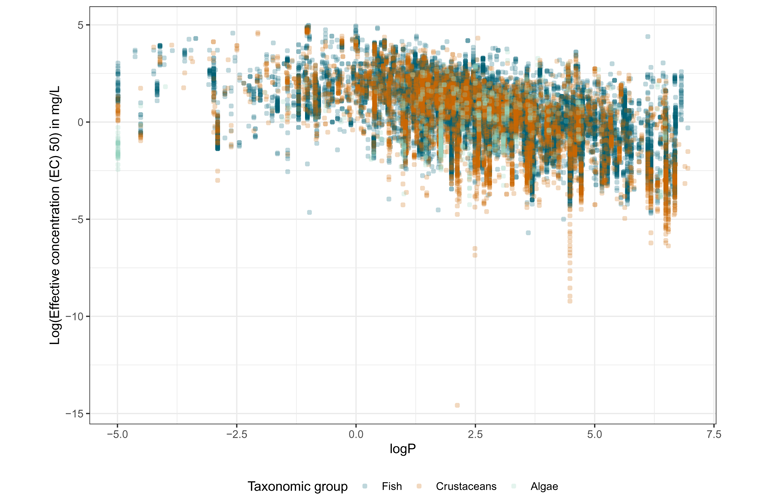
Relationship between logP and the effective concentration 50 (EC50, in *mg/L*) for the three taxonomic groups. Point transparency is set to 25 % to account for overplotting in areas with high point density.

**Figure 8.**
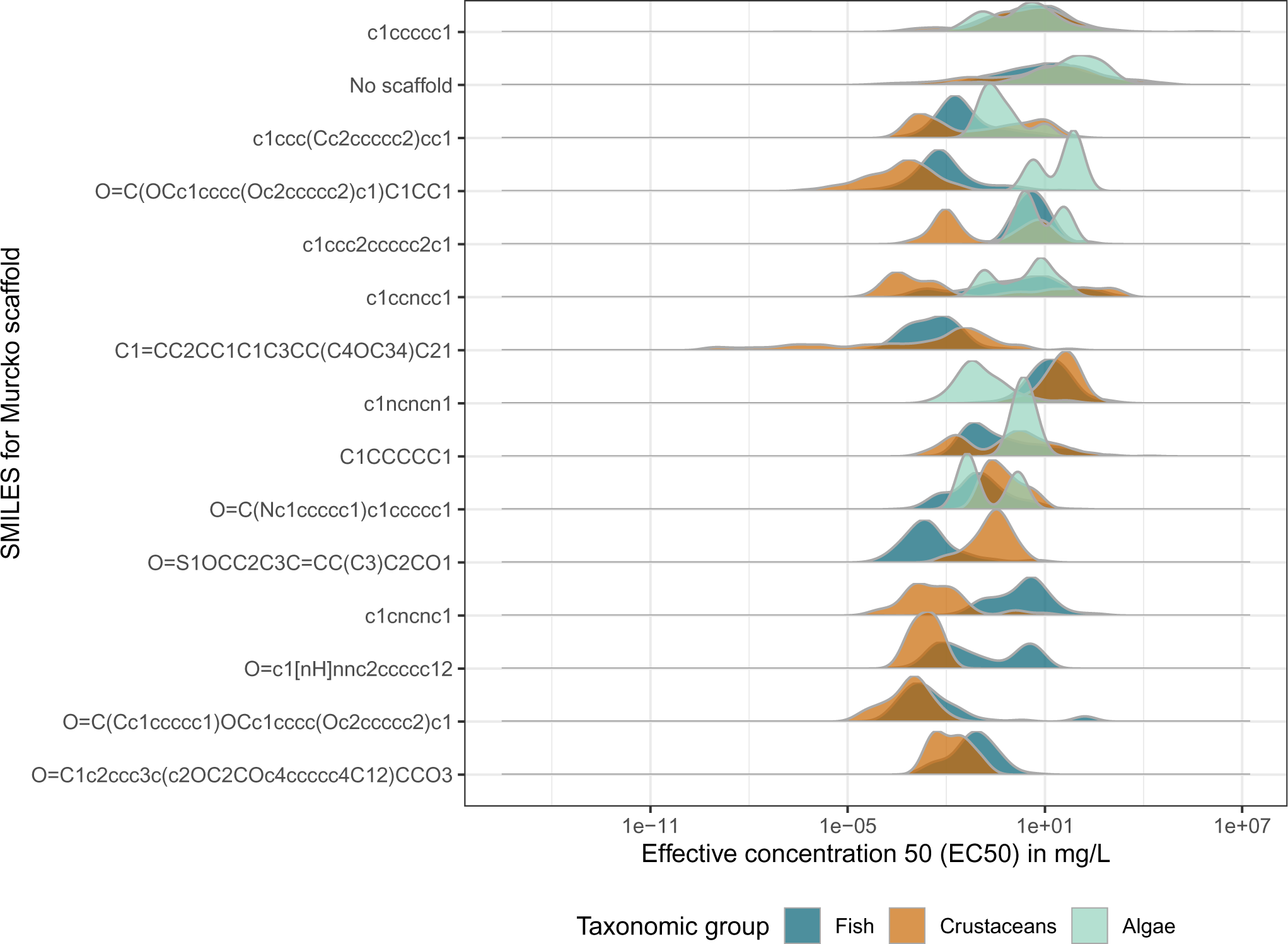
Effective concentration 50 (EC50, in *mg/L*) values for the top 15 Murcko scaffolds.

A relationship between logP and toxicity has been established for a number of chemicals, most commonly those with a Verhaar classification as non-polar narcotics (Verhaar category I) and polar narcotics (Verhaar category II)^63,64^. While not linear across the whole spectrum of logP values, this relationship forms the basis of many QSAR models^63^. These models then are limited in their applicability domain for the logP range for which this relationship has been established. For *ADORE*, the relationship of logP and toxicity is shown in Figure 7 for the three taxonomic groups. While the relationship is visible, we highlight that this relationship is noisy (indicated by the range of data points per logP value) and the LC50 cannot be only explained through the logP. The majority of chemicals in *ADORE* could not be assigned to an informative Verhaar class with the QSAR toolbox^65^ (data not shown), which is in line with the findings of Ebbrell *et al.*^63^. They reported out of domain categorizations (Verhaar class V) for 31% of industrial chemicals in their dataset and as much as 70% of active pharmaceutical ingredients. Our goal with *ADORE* was to compile a broadly applicable dataset that can be used beyond the applicability domains of conventional QSARs and not only for specific modes of action. To reduce the potential for data leakage, we did not include any mode of action classifications in the dataset.

Additional information on the functional use categories is given in Annex Figure 16 as an overview of the most prevalent OECD functions and their non-harmonized counterpart. For 570 of the 2,408 chemicals in our dataset, we could determine a functional use. Biocides are by far the most common, making up 43%, followed by fragrances (11%) and solvents (9%). For the biocides, insecticides are most common (21% of all chemicals with a functional use category), followed by fungicides (13%, and therefore more common than fragrances and solvents), and herbicides (3%). This result is not surprising as an ecotoxicological dataset is inherently biased towards toxic compounds. The information on the most common functional use categories has to be considered with caution as no function use information is available for most of the data.

## Data splittings

We have a closer look at three splitting schemes representative for the three splitting categories in Figure 9: *totally random* (A) where data leakage is expected, *by occurrence of chemical compounds* (B), and *by Murcko scaffold* (C). For each splitting, we compare the distributions for toxicity (as EC50), molecular weight, and logP for the training and test sets. *Totally random* splitting results in coinciding distributions of the three parameters across both training and test set. Splitting *by occurrence* and *by Murcko scaffold* produces EC50 distributions that cover a similar range but peaking at different EC50 values, which is more pronounced for the splitting *by Murcko scaffold*. The distributions for molecular weight and logP are less similar, but generally cover similar ranges. Therefore, predictions based on these splits originate from the same the toxicity range they were trained on. Intuitively, the model performance should be better when the toxicity range for the test set is included in the range that was contained in the training set. Exceptions can arise from activity cliffs, structurally similar compounds that differ strongly in biological activity^66,67^.

**Figure 9.**
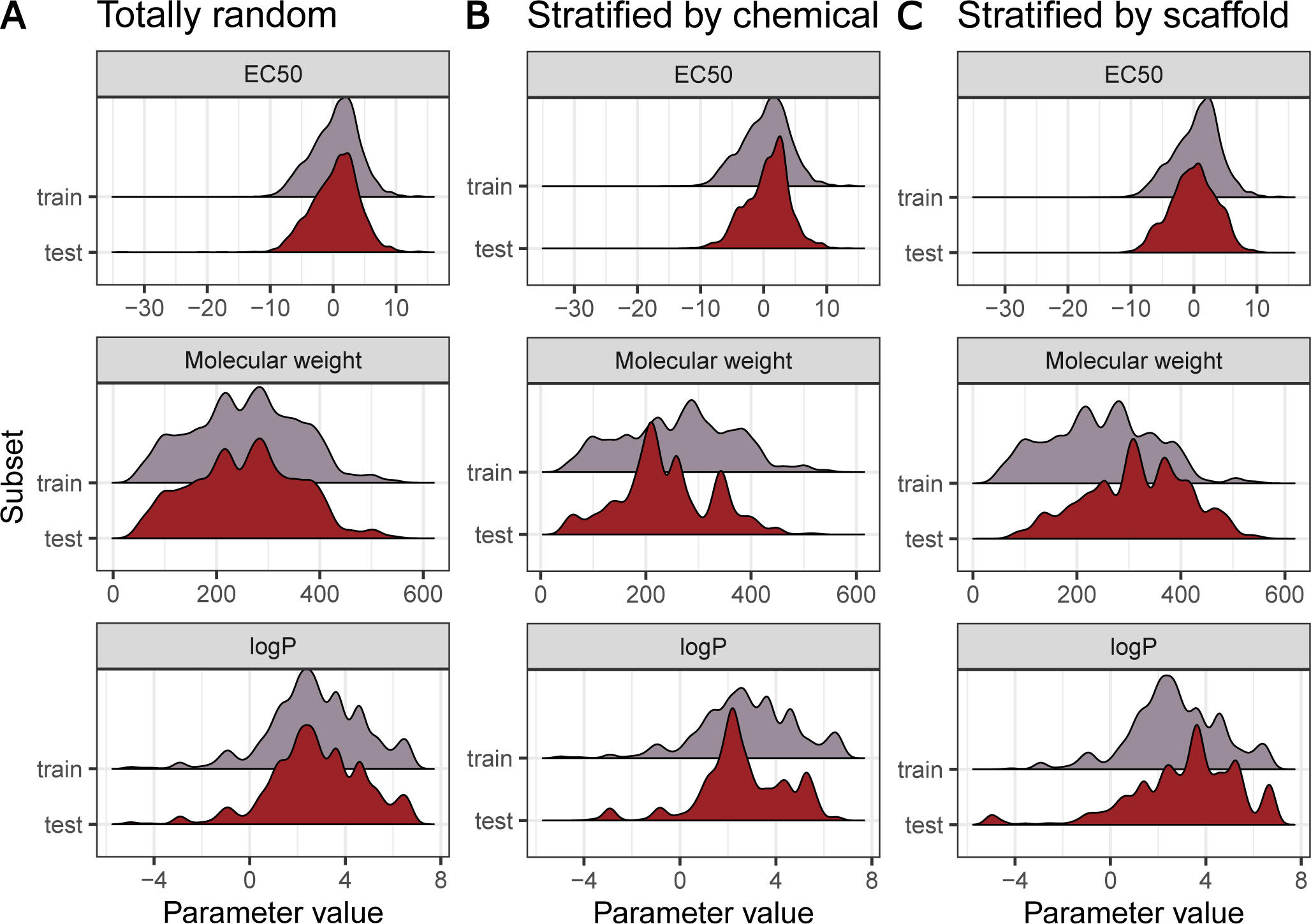
Overview of EC50 (log10-transformation of the mass concentration in *mg/L*), molecular weight (in *g/mol*), and logP for three data splittings: A: *Totally random*, B: *by occurrence of chemical compound*, and C: *by Murcko scaffold*.

Intuitively, splitting by scaffold (with the second largest compound group having “no scaffold” due to a lack of ring structures), presents a very challenging generalization task. During training, the model will only be presented with a rather heterogenous subset of chemicals that are very different from the chemicals in the other scaffold bins.

## Usage Notes

### Prediction Challenges

Here, we propose several tasks to be addressed with our dataset, so, ultimately, results can be compared between modeling studies based on this dataset. For each data subset, we provide the training and test sets as described in the corresponding “Methods” subsection. We also provide cross-validation folds if the splitting scheme is suitable for it. Model selection should be performed on the training set, and the test set should only be used once, at the end of the study, to assess the generalization performance of the model. The test set *must* not be used for model training^10^.

For all challenges, we suggest to split by occurrence of chemical compounds which prevents data leakage through the same chemicals being both in training and test sets. Splits by scaffolds are worth exploring to gain a better understanding of how they affect model outcome and whether they are suitable for splitting schemes. We strongly advise against the use of the *totally random* split as it will lead to overestimated model performances.

### Using the whole dataset

Ultimately, the goal of *in silico* methods is to replace *in vivo* experiments, at least with vertebrates. We propose two ways to approach this. The first option is a generalization to species, *i.e.,* train on crustaceans and algae, and test the models on fish (Table 3, a-CA2F-same and a-CA2F-diff). If it is possible to learn from invertebrate and algae *in vivo* data, which are considered less ethically problematic and more cost efficient, the number of tests on fish may be reduced in the future. This generalization to species can be performed either on the same chemicals (a-CA2F-same) or on different chemicals (a-CA2F-diff). The latter corresponds to a generalization to both species and chemicals, which presumably is the hardest challenge we propose. The second option is to train and test on all data of the three taxonomic groups to improve predictions for the species and chemicals available in the dataset (a-FCA2FCA).

### Within a taxonomic group

The three taxonomic groups might be too different to be modelled together. Alternatively, we propose prediction challenges within each taxonomic group (Table 3). These challenges may help to gain a better understanding of mortality within each taxonomic group and to determine whether the inclusion of species-related features helps to find explainable models with good prediction quality.

### On single species

Since most previous studies focused on single species, we also provide challenges for the most prominent species for each taxonomic group in our dataset (Table 3). For fish, we provide the top 3 species whereas for crustaceans and algae, we only provide the top species. For these challenges, species-specific features are not informative and need to be ignored.

## Limitations

We would like to address some limitations that concern the field of predictive ecotoxicology in general and our dataset in particular.

*In vivo* data is, by nature, highly variable and our core dataset spans several decades of experimental work (Annex Figure 10). During that time, several standards and test guidelines, like those from the ISO and OECD, were instituted to achieve higher reproducibility of experiments. Similarly, not all information desirable for modeling is contained in the original sources. Generally, the abundance of features comes with the caveat that not all parameters were recorded in all experiments, resulting in a fair amount of “Not recorded” (NR, NC) values. For some parameters we remove those entries, for others they make up the majority of entries and could therefore not be removed, *e.g.,* the organism life stage as shown in Annex Figure 11E. The inclusion of an appropriate control treatment is under most circumstances a requirement of the ECOTOX selection process^15^. Nonetheless, the encoding in the database includes the factors “inappropriate”, “unsatisfactory”, “not reported”, and “unspecified”. These encodings indicate for which entries the desirable control types, *i.e.,* negative, positive, and solvent control, were not included in the original study^68^). Nonetheless, certain data points are included by ECOTOX, when the use of controls can be deduced from the original source. Here, we decided to trust the selection criteria for the ECOTOX database, but advise against using control type as modeling feature (Annex Table 6).

Furthermore, the toxicity of different compounds, especially metals, is highly dependent on the medium composition and medium properties. This behavior is very complex and hard to predict, but to a degree interlinked with the acid dissociation constant pKa^69^. Therefore, we included this parameter such that its impact on model performance can be studied. Concurrently, isomerism and 3D structures of molecules are not captured by the molecular representations used here.

Ideally, chemical properties, molecular representations, and species-related information would fully explain all toxicity outcomes by covering every physical, chemical, and biological dimension of an experiment. This, however is not the case with the features currently available. Mortality in itself is a highly unspecific endpoint that can be caused through a plethora of different mechanisms, particularly when considering the differing biology of the studied organism types fish, crustaceans, and algae. Cheminformatics use cases following a more mechanistic focus, *e.g.,* modeling the interaction of a molecule with a specific molecular targets such as triggering a molecular event/receptor response/gene expression change;^70^, likely have a higher benefit from molecular representations, since there is an immediate cause-and-effect-relationship between the molecular structure and the molecular target at hand. With predictive ecotoxicology still in its infancy, this dataset hopefully fosters exploration and a better understanding of what kind of data is helpful or lacking to further progress the field. Accordingly, our goal is to help the field of predictive ecotoxicology progress by enabling reproducibility and comparability across studies based on our dataset.

## Model implementation

Here, we highlight best practices and common pitfalls to be considered when modeling with the *ADORE* dataset. Recently, several researchers have addressed sources of reproducibility issues in ML-based research^10,71,72^. Checklists on how to circumvent these pitfalls by following best practices regarding model implementation and reporting have been published^27,73,74^. Accordingly, we urge researchers to adhere to these best practices, to acquaint themselves with common issues, and to strive to serve as positive examples when implementing and reporting their models, results and conclusions. As previously discussed, the dataset contains many repeated experiments, increasing the likelihood of data leakage and, as a consequence, overestimated model performances^10^. To avoid this and to ensure comparability of studies based on *ADORE*, we urge users to adhere to the exact splittings we provide with the different challenges. This precaution is in line with the three checkpoints on data leakage by Kapoor et al. (2023)^73^: 1. separation of training and test data is maintained throughout the process and the test set is only used at the very end, *e.g.,* ensure to preprocess features only on the training data; 2. be mindful of dependencies or duplicates, *i.e.,* do not use the totally random split; 3. feature legitimacy, *i.e.,* do not use features that are proxies for the target response (mode of action, labels of the Globally Harmonized System of Classification and Labelling of Chemicals (GHS) system or other hazard classifications, *etc.*)^73^. In the glossary, we indicate which features are suitable for modeling.

## Code availability

The code used to load and process the input data and generate the output dataset was created and run in Python 3.9 and is made available on https://renkulab.io/gitlab/mltox/adore. The repository contains code on how to load the data, prepare it for modeling, *e.g.,* create one-hot and multi-hot-encodings for categorical features, and apply the train-test-split for 5-fold cross-validation. A good starting point are the files in the folder *scripts* for random forests (*14_analysis_regression_rf.py* and *34_analysis_regression_test_rf.py*).

## Acknowledgements

This work was made possible through the SDSC grant “Enhancing Toxicological Testing through Machine Learning” (project NoC20-04) and partly carried out in the framework of the European Partnership for the Assessment of Risks from Chemicals (PARC) and has received funding from the European Union’s Horizon Europe research and innovation programme under Grant Agreement No 101057014. Views and opinions expressed are however those of the author(s) only and do not necessarily reflect those of the European Union or the Health and Digital Executive Agency. Neither the European Union nor the granting authority can be held responsible for them. Some figures were created with BioRender.

## Author contributions statement

L.G. and C.S.: Conceptualization, Software, Validation, Formal analysis, Investigation, Data curation, Visualization, Writing - Original Draft, Writing - Review & Editing. F.P.C, K.S., M.B.J.: Conceptualization, Supervision, Funding acquisition, Writing - Review & Editing. All authors reviewed and approved the manuscript.

## A Glossary

**Table 6.**
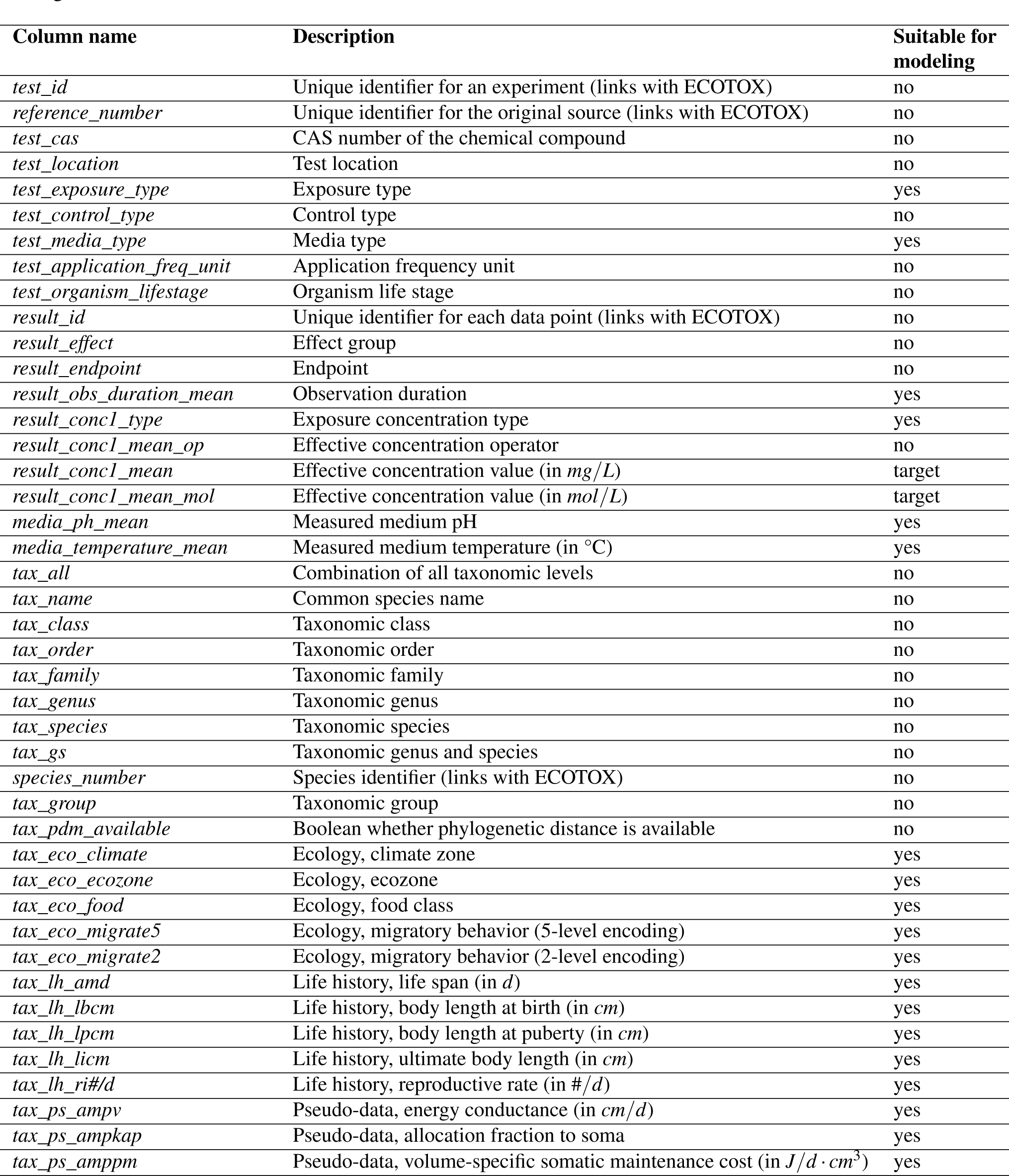
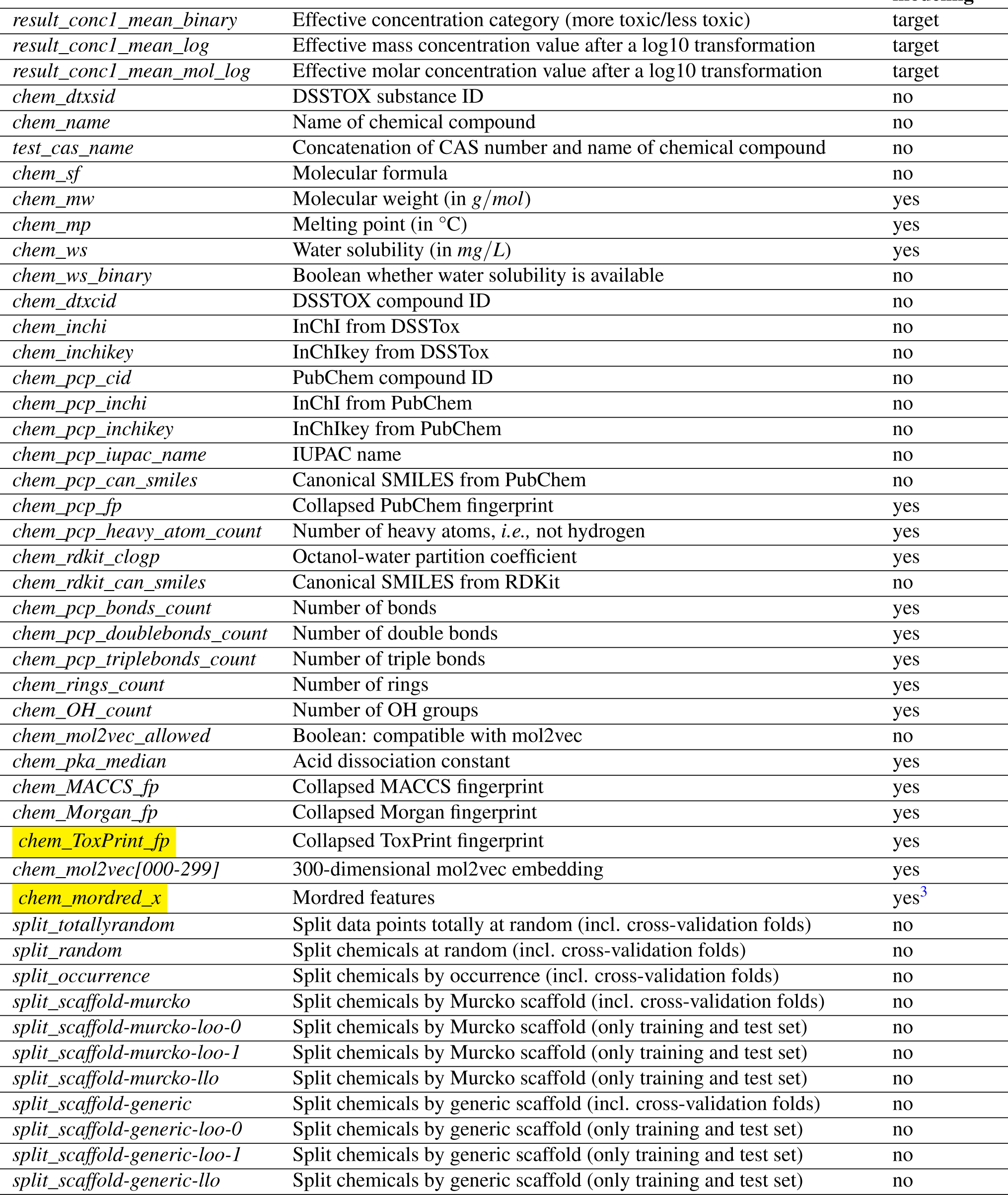
A glossary of all columns in the dataset. The prefix of the column name identifies the feature category or source file, *e.g.,* column names starting with “test” originate from the ECOTOX tests file. The dataset contains modeling features as well as additional information. Only the features labelled “suitable for modeling” should be used for modeling. The others *must not* be used. Effective concentrations are labelled as targets. Only one target should be used and the others *must not* be used as modeling feature.

### B ECOTOX data

Definitions for ECOTOX effect groups as given in the ECOTOX term appendix:

- Mortality (MOR): Measurements and endpoints where the cause of death is by direct action of the chemical.
- Physiology (PHY)/Intoxication (ITX): Measurements and endpoints regarding basic activity in cells and tissues of plants or animals; includes four effect groups - injury, immunity, intoxication and general physiological response.
- Growth (GRO): Category encompasses measures of weight and length, and includes effects on development, growth and morphology.
- Population (POP): Measurements and endpoints relating to a group of organisms or plants of the same species occupying the same area at a given time.

#### Toxicity categories

The toxicity intervals given in Table 7 and shown for the chemicals in our dataset in Figure 3 are in accordance with EPA.

**Table 7.** EC50 intervals for the binary and multi-class toxicity classification used in Figure 3.

| EC50 interval (mg/L) | Binary classification | Multi-class classification |
| --- | --- | --- |
| $(-\infty, 10^{-1}]$ | more toxic | very highly toxic |
| $(10^{-1}, 10^0]$ | more toxic | highly toxic |
| $(10^0, 10^1]$ | less toxic | moderately toxic |
| $(10^1, 10^2]$ | less toxic | slightly toxic |
| $(10^2, +\infty)$ | less toxic | non-toxic |

**Figure 10.**
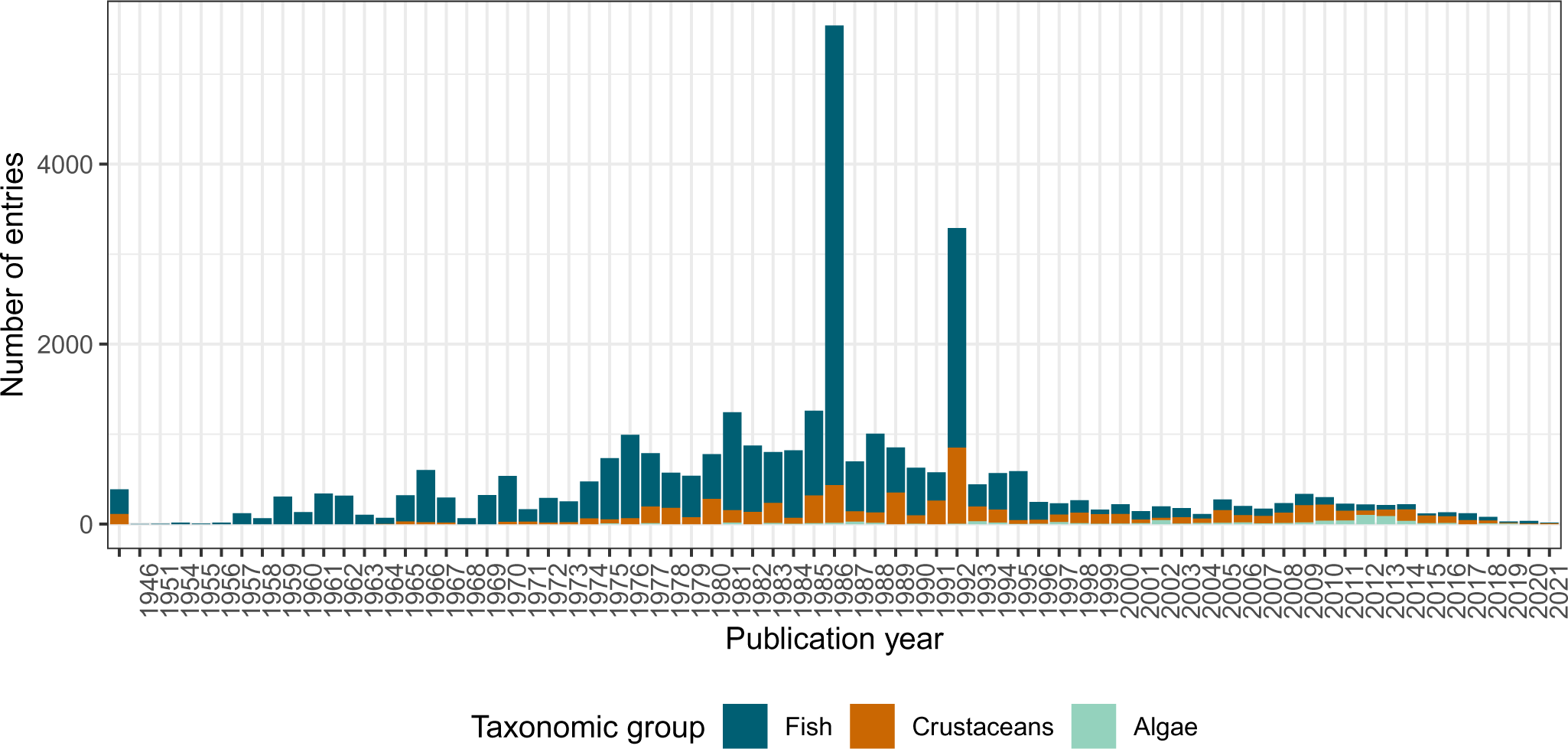
Chronology of the references related to the different taxonomic groups added to ECOTOX over time. The publication year refers to the year the original study was published. It is noteworthy that some references are not singular scientific studies, but entail whole databases, such as the US EPA “Pesticide Ecotoxicity Database (Formerly: Environmental Effects Database (EEDB))”, added in 1992.

#### Reference chronology

### B.1 Experimental properties

**Figure 11.**
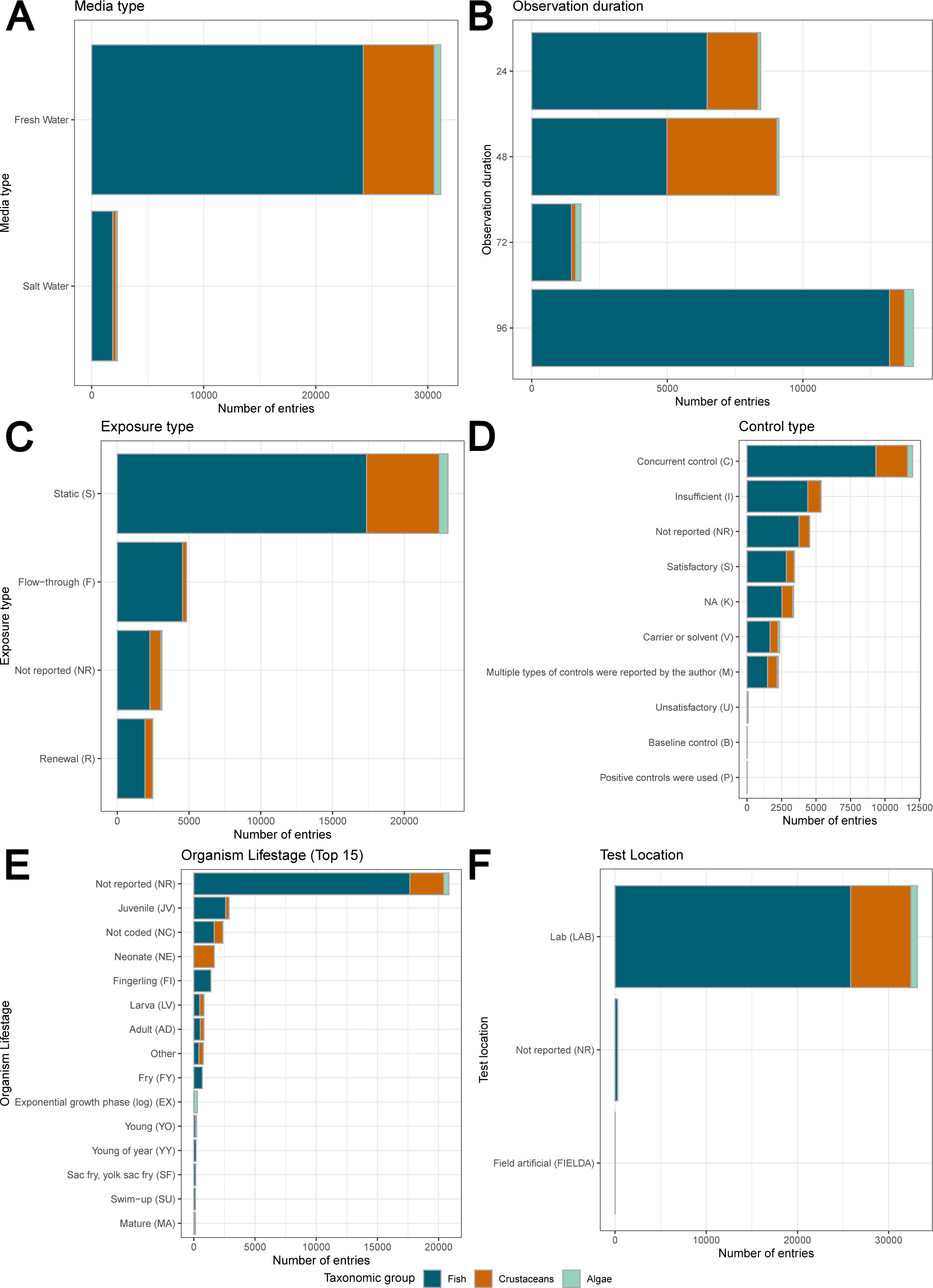
Overview of the most common values for experimental properties parameters. A: Media type; B: Observation duration; B: Exposure type; D: Control type; D: Organism life stage (only the top 15 of 48 total, less-abundant life stages are combined into “other”); F: Test location.

### C Taxonomic data

#### C.1 Overview

**Figure 12.**
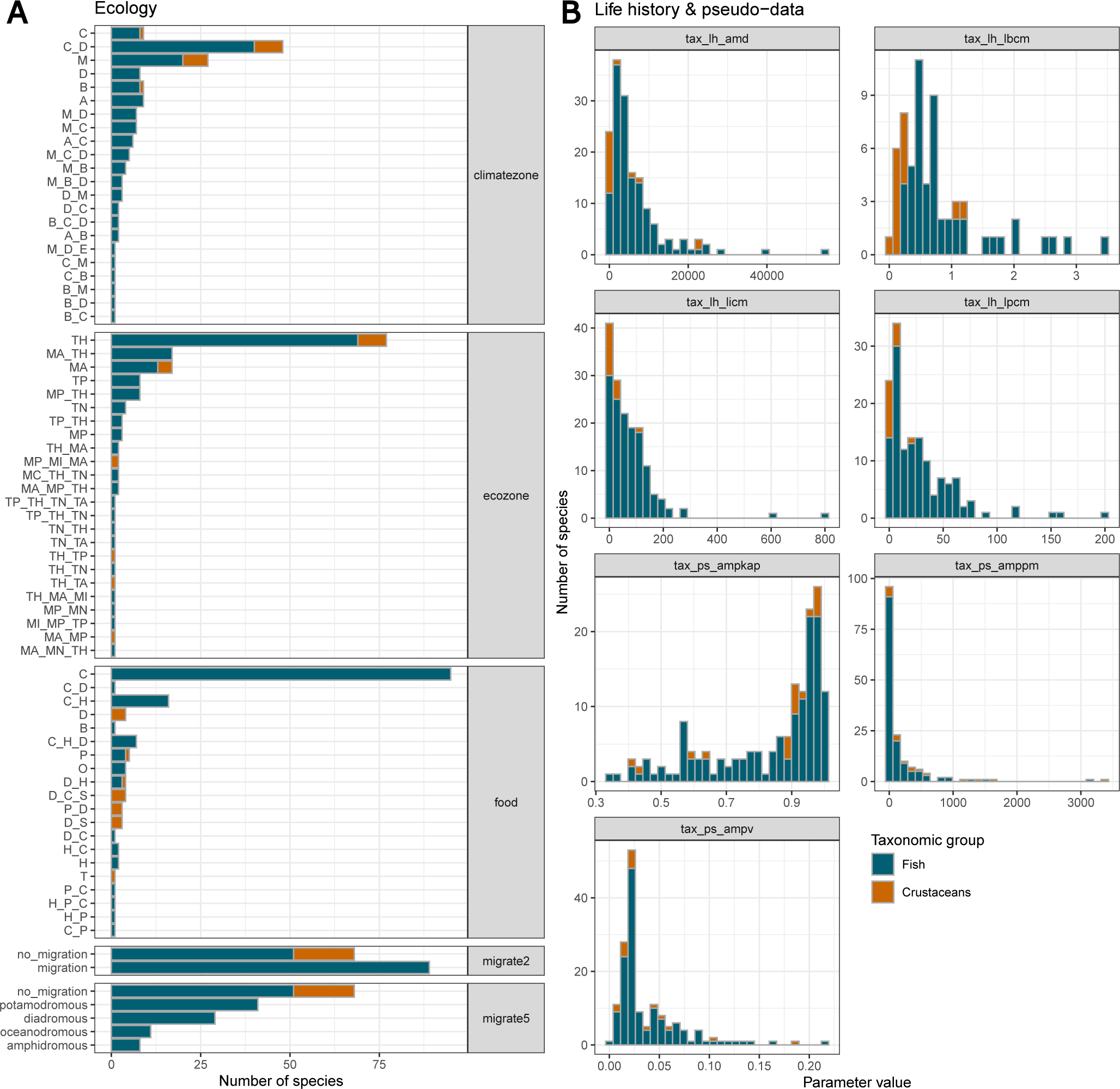
A: Overview of the taxonomic properties for ecology. B: Overview for life history and pseudo-data parameter values. The encodings for the different levels are given in Annex Tables 8-11.

#### C.2 Ecological data encoding

Here, we provide the encodings for the ecological data. For some parameters, we had to reduce the complexity of the encodings. More detailed descriptions of the parameters and the original encoding can be found in the Add my Pet collection

##### C.2.1 Climate zone

**Table 8.** Climate zone definitions and encodings.

| Encoding | Definition |
| --- | --- |
| A | Tropical (megathermal) climates: every month of the year with an average temperature of 18 °C or higher, with significant precipitation |
| B | Dry (arid and semi-arid) climates: little precipitation |
| C | Temperate (mesothermal) climates: the coldest month averaging between 0 and 18 °C and at least one month averaging above 10 °C |
| D | Continental (microthermal) climates: at least one month averaging below -3 °C and at least one month averaging above 10 °C |
| E | Polar and alpine (montane) climates: every month of the year with an average temperature below 10 °C |
| M | Marine climates |

##### C.2.2 Ecozone

**Table 9.** Ecozone definitions and encodings.

| Encoding | Definition |
| --- | --- |
| MA | Atlantic ocean |
| MC | Circumglobal oceans |
| MI | Indian ocean |
| MN | Arctic ocean |
| MP | Pacific ocean |
| MS | Southern ocean |
| TA | Australasia (including New Guinea, New Zealand) |
| TH | Holarctic |
| TN | Neotropic (including Central America, and the Caribbean) |
| TP | Paleotropic |

##### C.2.3 Migratory behavior

**Table 10.** Migratory definitions and encodings for 2 and 5 levels.

| 2-level encoding | 5-level encoding | Definition |
| --- | --- | --- |
| no_migration | no_migration | Do not migrate |
| migration | amphidromous | Migrate from fresh water to the sea, or vice versa, but not for breeding |
| migration | diadromous | Migrate between the sea and fresh water |
| migration | oceanodromous | Live and migrate wholly in the sea |
| migration | potamodromous | Live and migrate wholly within fresh water |

##### C.2.4 Food

**Table 11.** Food definitions and encodings.

| Encoding | Definition |
| --- | --- |
| B | Bacterivore (including micro-organisms) |
| C | Carnivore (living animals) |
| D | Detrivore (bacteria, small fungi, organic matter) |
| H | Herbivore (plants) |
| O | Omnivore (plants/animals/fungi) |
| P | Planktivore (small aquatic organisms, macro-movement controlled by flow, not by swimming) |
| S | Scavenger (dead animals) |
| T | Parasitic (animal tissue) |

### D Chemical data

#### D.1 Tanimoto similarity

**Figure 13.**
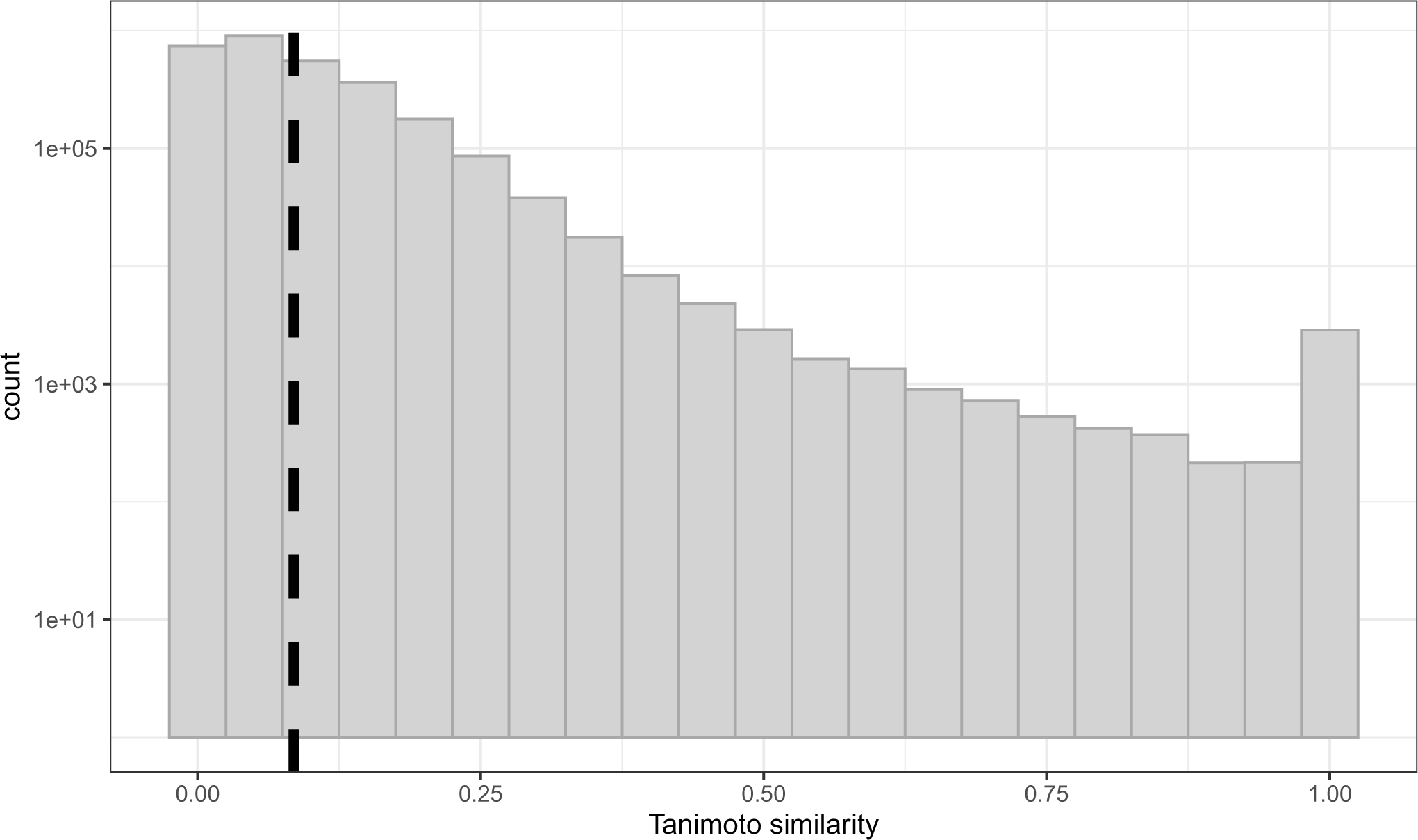
Histogram of the Tanimoto similarities for the chemicals in our dataset, ranging from 0 (dissimmilar) to 1 (equal). The dashed vertical line indicates the mean similarity of 0.085.

#### D.2 Chemical ontology

**Figure 14.**
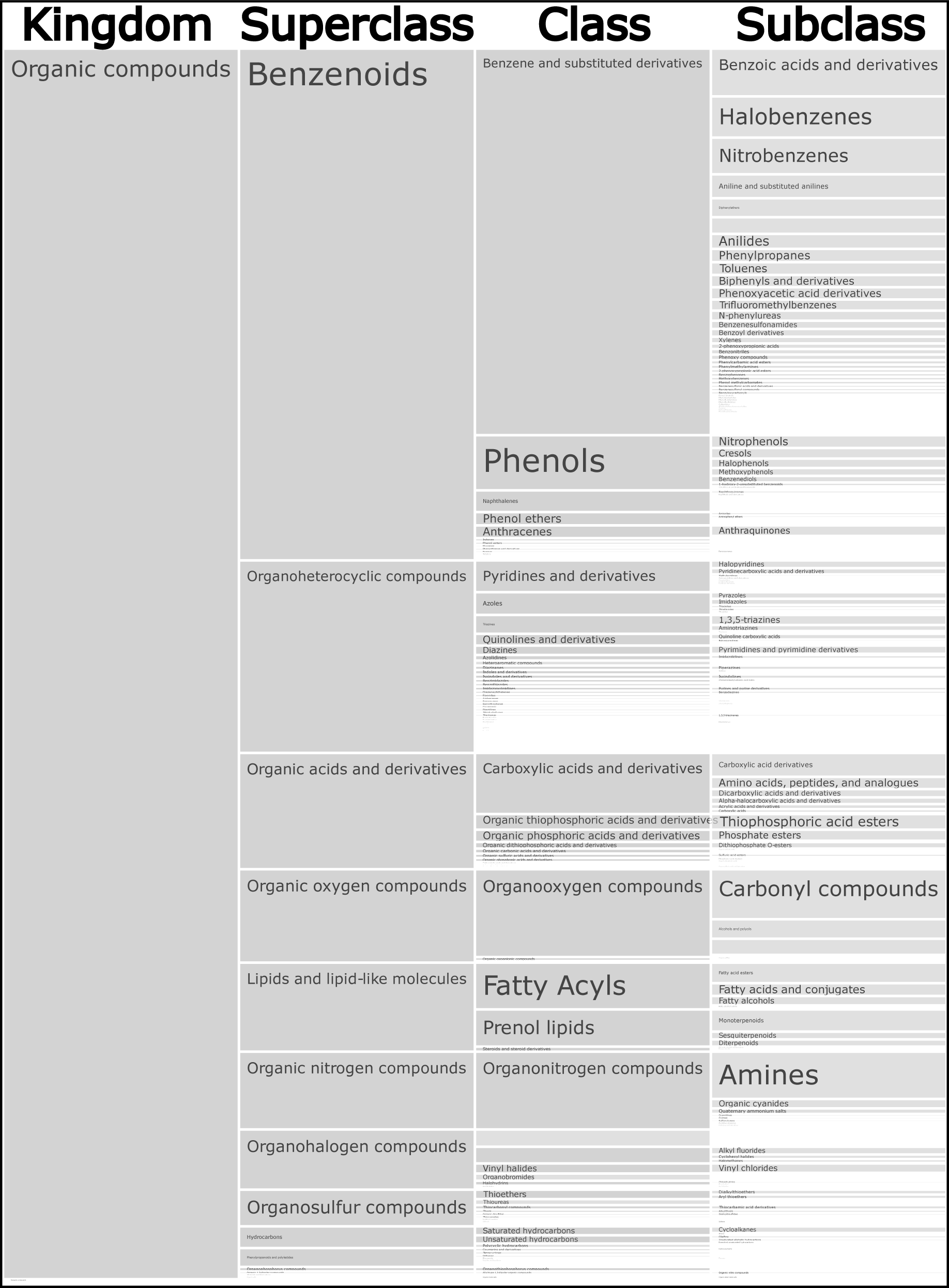
Icicle chart representing the chemical ontology for 2,250 of the 2,408 chemicals in our dataset, with the hierarchical levels kingdom, superclass, class, and subclass.

#### D.3 Chemical properties

**Figure 15.**
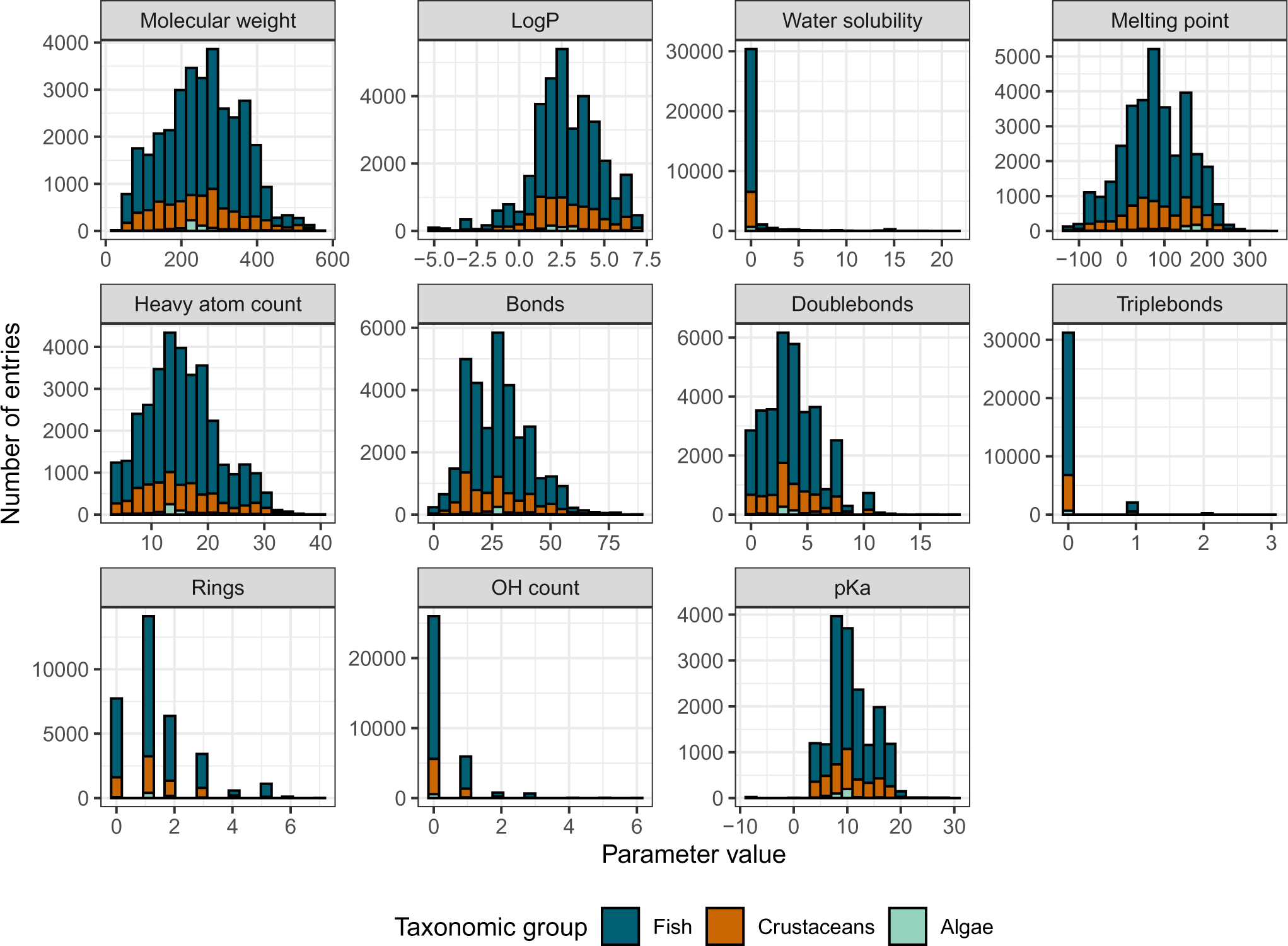
Overview of the distributions of the chemical properties represented in our dataset. See Table 5 for more details.

#### D.4 Functional use categories

**Figure 16.**
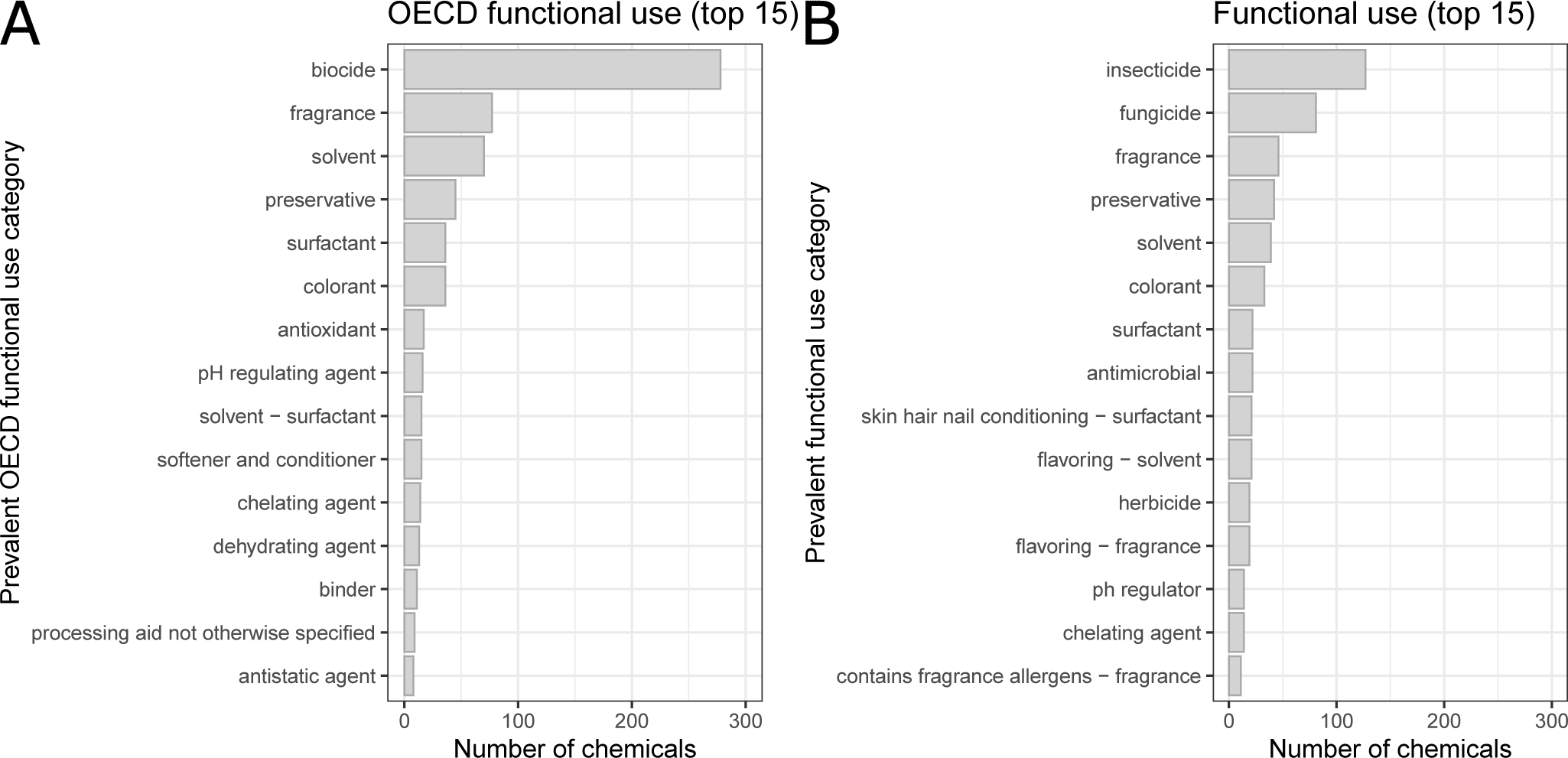
Prevalent functional use categories for 570 of the 2,408 chemicals in our dataset. Most chemicals have several reported uses of which we select the prevalent ones, *i.e.,* those reported in at least 20% of the cases for each chemical. A: Internationally harmonized OECD functional uses, B: non-harmonized functional uses distinguish between the different biocides.

#### D.5 Molecular scaffolds

**Figure 17.**
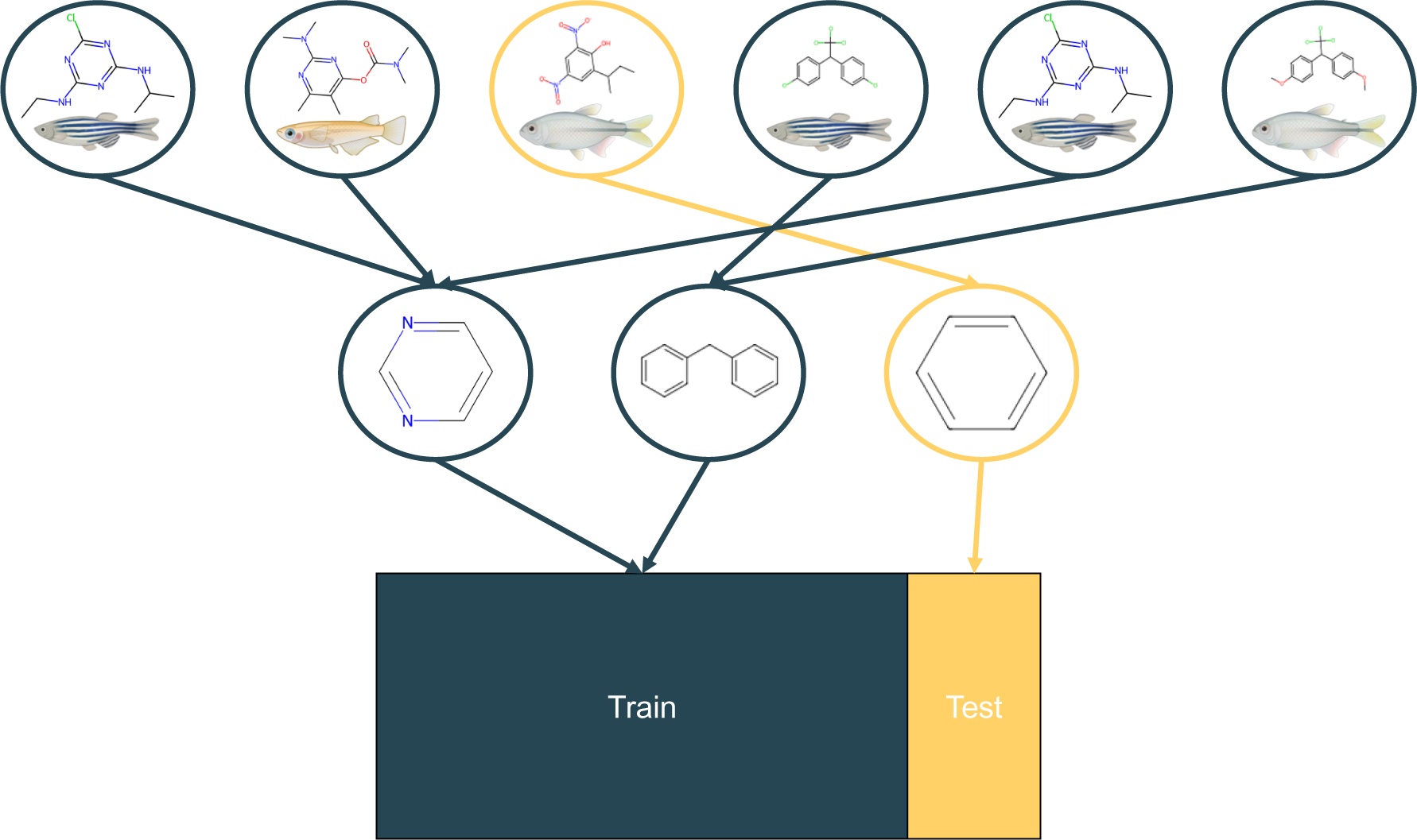
Schematic visualization of stratified data splitting according to the molecular scaffold. In the first step, all molecules that share a molecular scaffold are grouped together. Then, these groups are distributed into a training and a test set. This procedure reduces data leakage by ensuring that similar chemicals - defined by a common molecular scaffold - do not occur both in the train and test set.

**Table 12.**
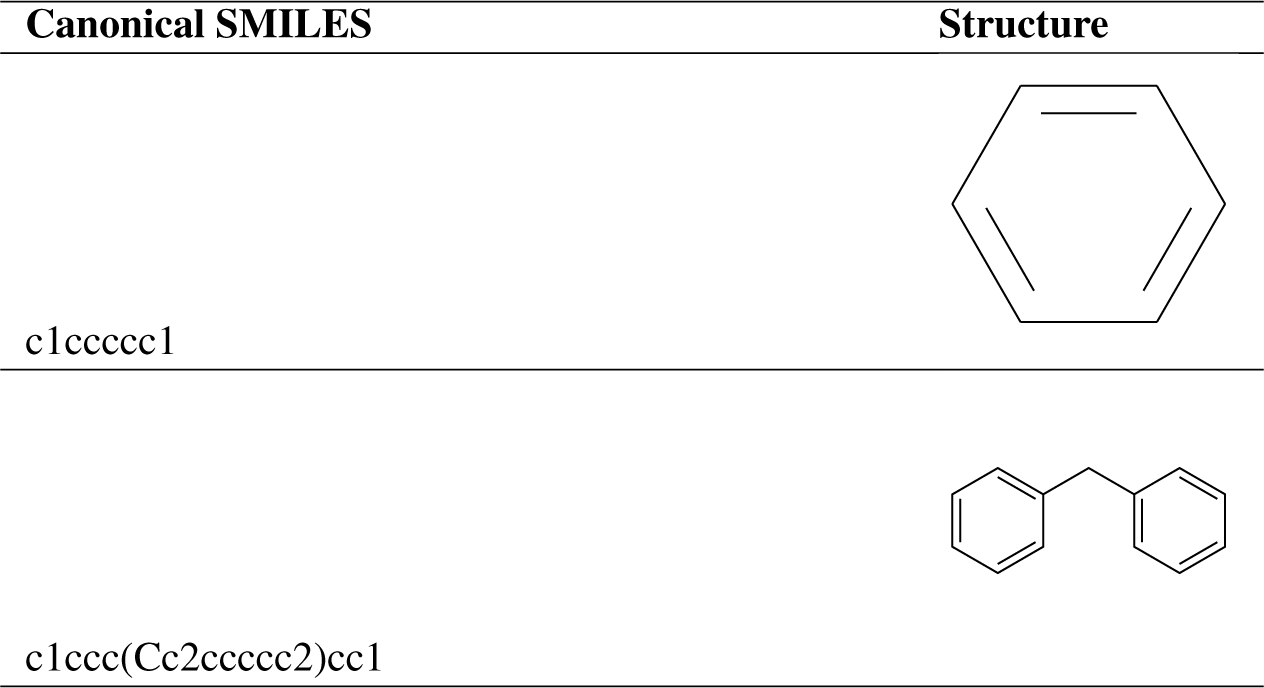
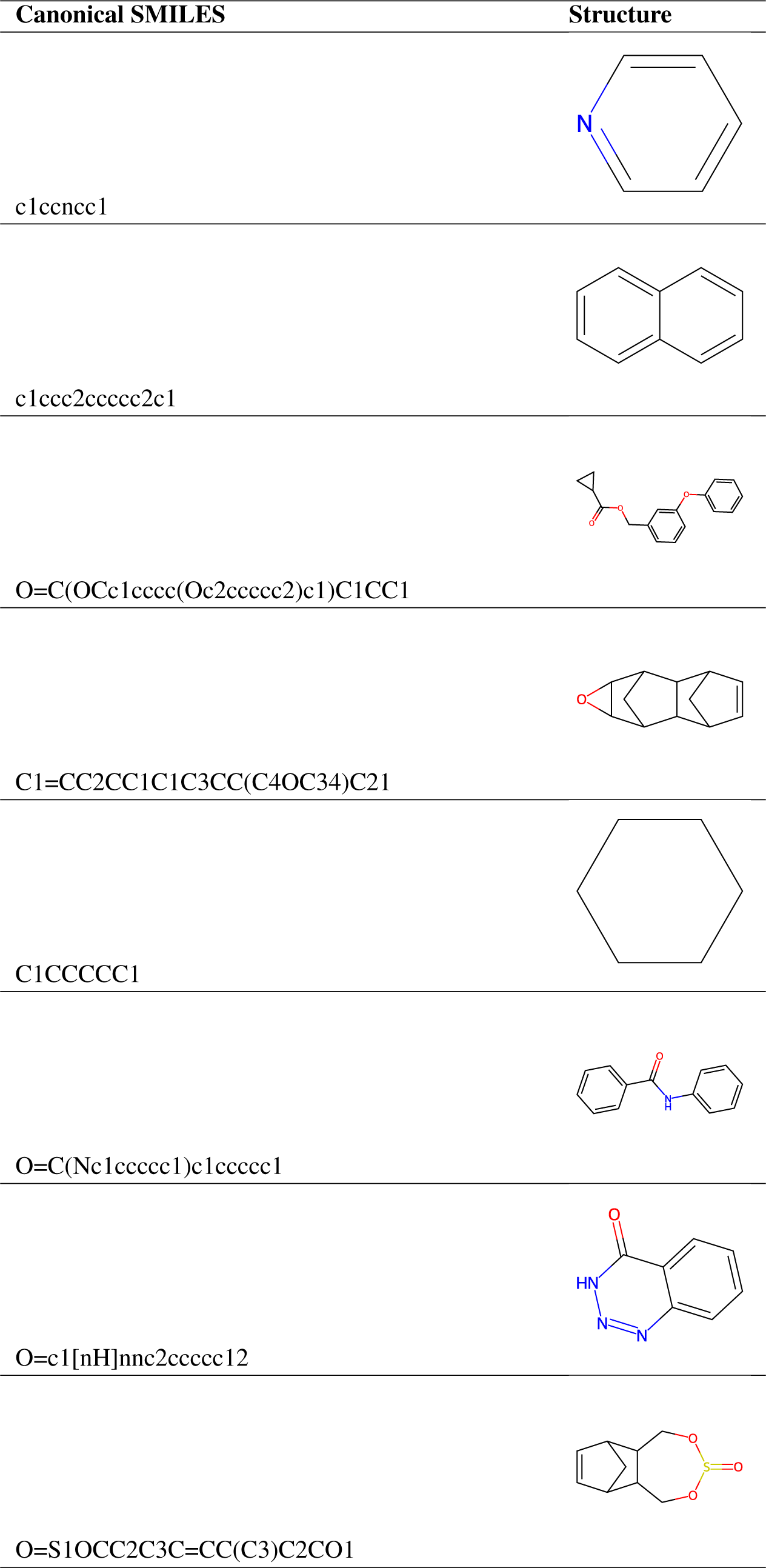
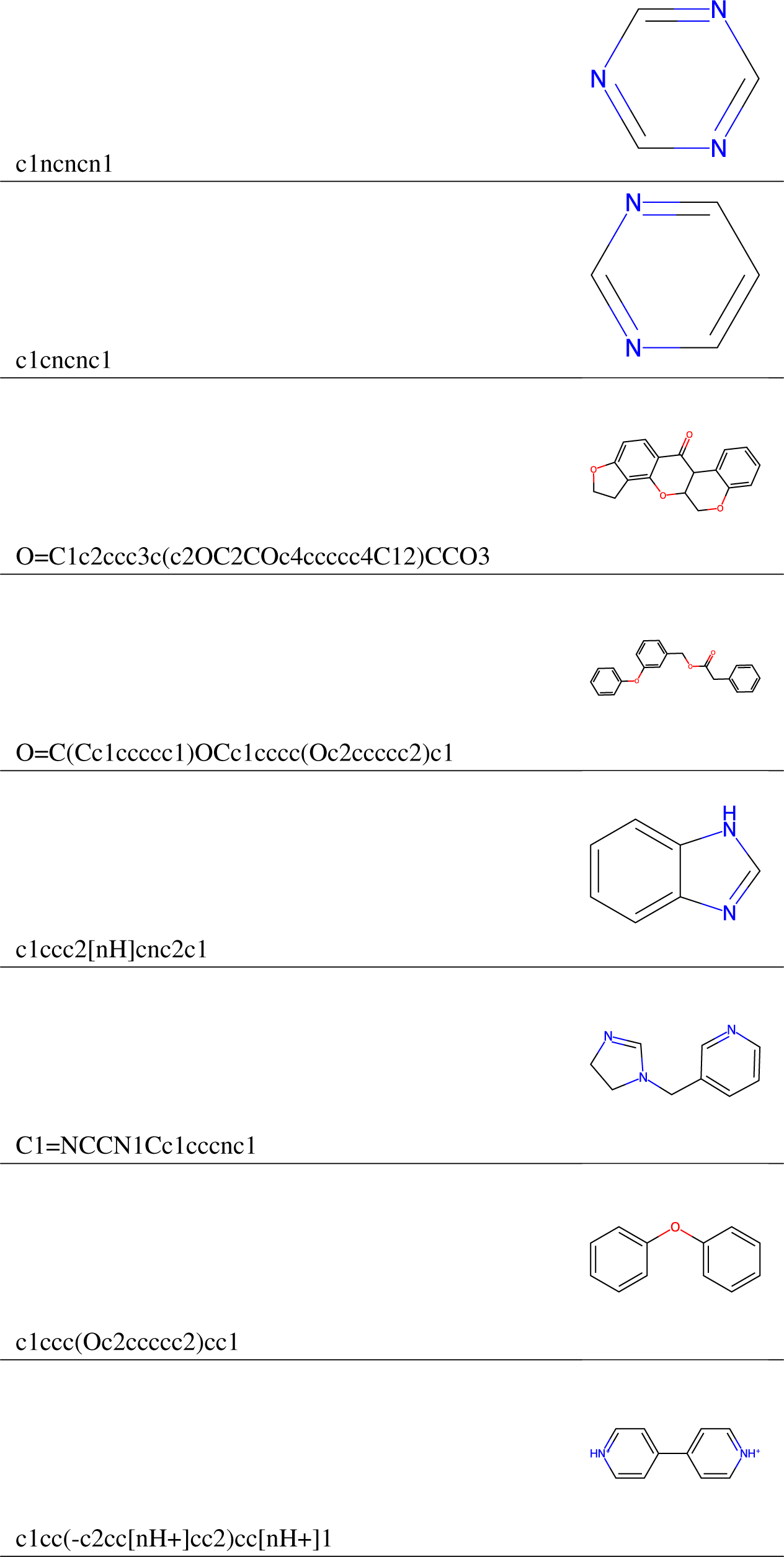
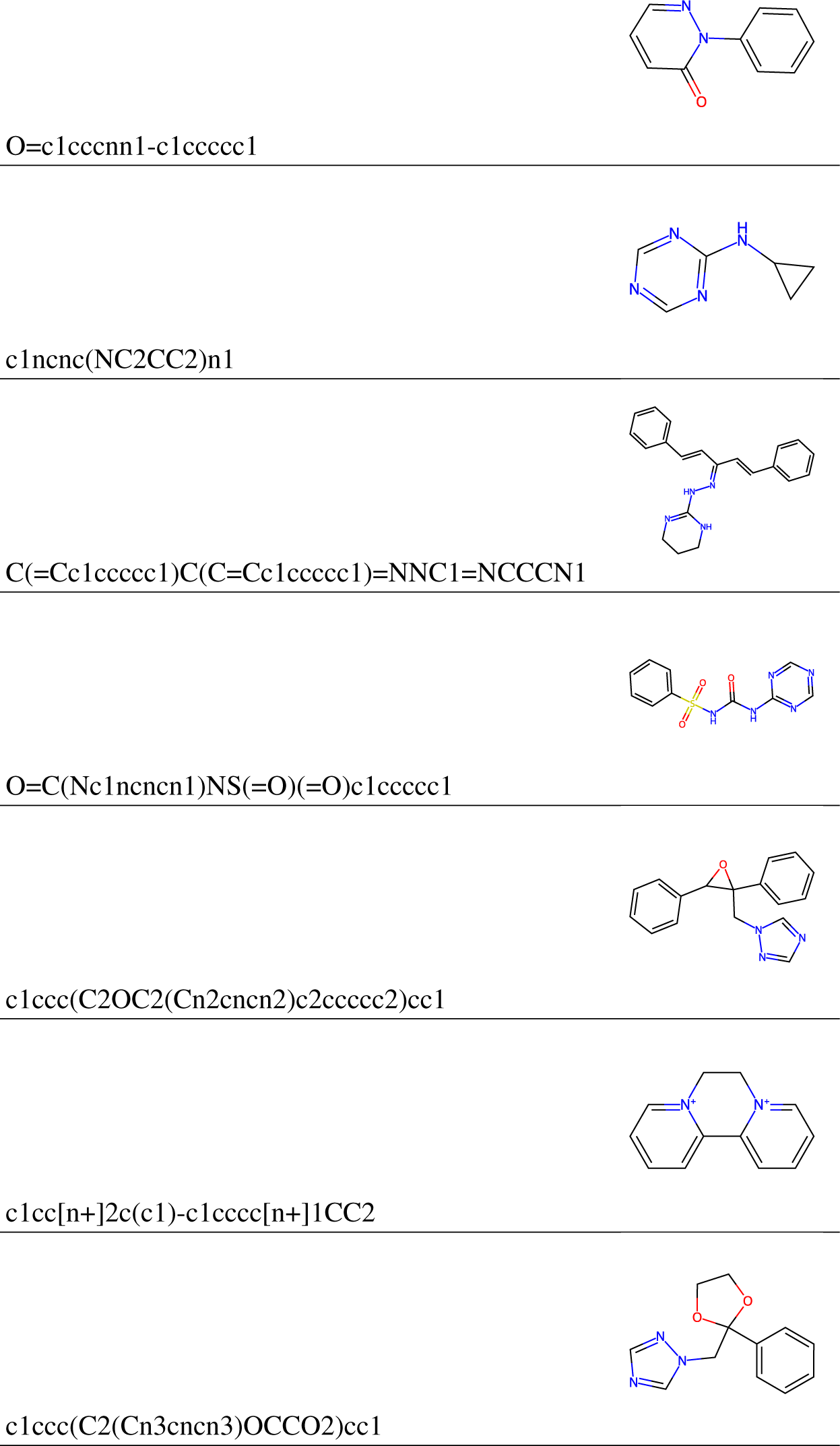
Canonical SMILES and chemical structures of the most common Murcko scaffolds. We do not include “no scaffold” here.

## Footnotes

1 Static (S, “Toxicity tests with aquatic organisms in which no flow of test solution occurs; solutions may remain unchanged throughout the duration of the test”), flow through (F, “continuous or very frequent passage of fresh test solution through a test chamber with no recycling”), renewal (R, “A test without continuous flow of solution, but with occasional renewal of test solutions after prolonged periods, e.g., 24 hours”), and not reported aquatic experiments (NR). These definitions are from the ECOTOX term appendix).

2 See these links for the definition of each MACCS bit and PubChem bit. Additionally, we added bit description files to the repository.

3 We advice modelers to investigate these features, e.g., with a correlation analysis, before adding them to the model.

